# Odor motion sensing enables complex plume navigation

**DOI:** 10.1101/2021.09.29.462473

**Authors:** Nirag Kadakia, Mahmut Demir, Brenden T. Michaelis, Matthew A. Reidenbach, Damon A. Clark, Thierry Emonet

## Abstract

Studies dating back a century (Flügge, 1934) have stressed the critical role of the wind as the primary directional cue in odor plume navigation. Here, we show that *Drosophila* shape their navigational decisions using a second directional cue – *the direction of motion of odors –* which they detect from the temporal correlations of the odor signal between their two antennae. Using a high-resolution virtual reality paradigm to deliver spatiotemporally complex fictive odors to freely-walking flies, we demonstrate that such odor direction sensing is computationally equivalent to motion detection algorithms underlying motion detection in vision. Simulations and theoretical analysis of turbulent plumes reveal that odor motion contains valuable directional information absent from the airflow; indeed, this information is used by both *Drosophila* and virtual agents to navigate naturalistic odor environments. The generality of our findings suggests that odor direction sensing is likely used throughout the animal kingdom, and could significantly improve olfactory robot navigation in harsh chemical environments.

## INTRODUCTION

Odor plumes in the wild are spatially complex and rapidly fluctuating structures carried by turbulent airflows (Riffell et al., 2008). Odors arrive in bursts of high concentration interrupted by periods of undetectable signal (Murlis et al., 1992; Murlis et al., 2000), and the temporal statistics of these odor encounters can vary by orders of magnitude (Celani et al., 2014). To successfully navigate odor plumes in search of food and mates, insects must extract and integrate multiple features of the odor signal, including the odor encounters’ intensity (Alvarez-Salvado et al., 2018; Pang et al., 2018), spatial distribution (Jung et al., 2015; Tao et al., 2020), and temporal aspects such as timing (Mafra-Neto and Cardé, 1994; van Breugel and Dickinson, 2014), duration (Alvarez-Salvado et al., 2018), and frequency (Demir et al., 2020; Jayaram et al., 2021; Kanzaki et al., 1992; Mafra-Neto and Cardé, 1994; Vickers and Baker, 1994). Effective plume navigation requires balancing these multiple streams of olfactory information and integrating them with other sensory inputs including visual and mechanosensory cues (Budick et al., 2007; Suver et al., 2019; van Breugel and Dickinson, 2014).

Like many animals, insects sense odors using two spatially separated sensors – their antennae which provides an information stream whose role in navigation still remains unclear. Indeed, *Drosophila* can detect inter-antennal concentration differences, and use them to navigate simple plumes such as static ribbons, where gradients are resolvable and informative (Duistermars et al., 2009; Gaudry et al., 2013). But the relevance of bilateral sensing for natural plume navigation is less clear, since odor gradients in turbulent flows fluctuate rapidly and do not reliably point toward the source (Alvarez-Salvado et al., 2018). Assessing whether insects use these gradients in complex plumes would require imaging odor signals in real-time during navigation, which was done for the first time only recently (Demir et al., 2020). While theoretical studies have suggested that gradients may be informative in near-surface turbulent plumes (Boie et al., 2018), this is not yet supported by observations (Alvarez-Salvado et al., 2018).

Here, we reveal a distinct role for bilateral sensing: detecting the direction of motion of odor signals. A waft of odor, such as a thin odor filament, passing laterally over an insect hits the two antennae sequentially; the filament’s direction of motion could in principle be inferred by resolving differences in firing rate between the antennae over time. Indeed, by reanalyzing data from an experiment in which odor plumes were measured simultaneously with fly behavior (Demir et al., 2020), we find a significant correlation between fly turning and odor motion direction. To investigate causality, we develop an optogenetic approach to deliver fictive odor signals with high temporal and spatial precision, and completely divorced from wind, to freely-walking *Drosophila*. In this setup, flies reliably turn against the direction of fictive odors, even in the absence of wind fly turning responses are odor *direction selective*. Leveraging stimuli from experiments exploring direction selectivity in the fly eye (Salazar-Gatzimas et al., 2016), we find that odor direction selectivity is consistent with elementary correlation-based algorithms underlying visual motion detection (Hassenstein and Reichardt, 1956), revealing the generality of these computations across sensory modalities. Naively, since odors are transported by the wind, odor motion and wind motion could be considered redundant directional cues. Instead, we find that odor direction sensing integrates with wind-driven responses in a mostly additive manner, and we show, using simulations of complex plumes, that odor motion contains valuable directional information absent in the airflow. To demonstrate the utility of odor direction sensing in a goal-directed task, we delivered complex fictive odor plumes and assessed flies’ ability to localize the source. Selectively perturbing odor direction, while leaving all other aspects of the plume and airflow unaltered, significantly degrades flies’ navigational performance. Finally, we show that complex plume navigation by virtual agents *in silico* is significantly enhanced by odor direction sensing, suggesting improvements in the design of olfactory robots. Our work reveals a key information stream for natural plume navigation, and suggests a valuable role for spatiotemporal sensing in environments which lack reliable odor gradients.

## RESULTS

### Flies respond direction selectively to odor motion in the absence of wind

To investigate if flies sense and react to odor direction, we first re-analyzed a dataset of walking *Drosophila* navigating a complex, visualizable odor plume whose odor statistics resemble those in turbulent flows (Demir et al., 2020) (Fig. 1a). In this plume, gradients can be randomly oriented relative to the source, and often differ substantially from the odor direction (Fig. 1a; green and magenta vectors). Since the odor is visible, we can quantify the odor signal perceived during navigation, as well as infer the projections along the antennae of the odor gradient and of the odor motion direction (Fig. 1b and Supplementary Fig. 1), while simultaneously measuring fly behavior (Fig. 1b). Insects turn upwind when encountering odor signals (Alvarez-Salvado et al., 2018; Budick and Dickinson, 2006; Demir et al., 2020; van Breugel and Dickinson, 2014), which we verified for flies oriented slightly away from the upwind direction (blue and red curves in Fig. 1c). For flies already oriented upwind, there was no odor-elicited turning bias, nor any turning bias relative to the perceived odor gradient (Fig. 1d). However, in this case, fly turning correlated significantly with odor *direction* (Fig. 1e), suggesting that flies use directional odor cues when directional information from the wind is minimized.

**Figure 1.**
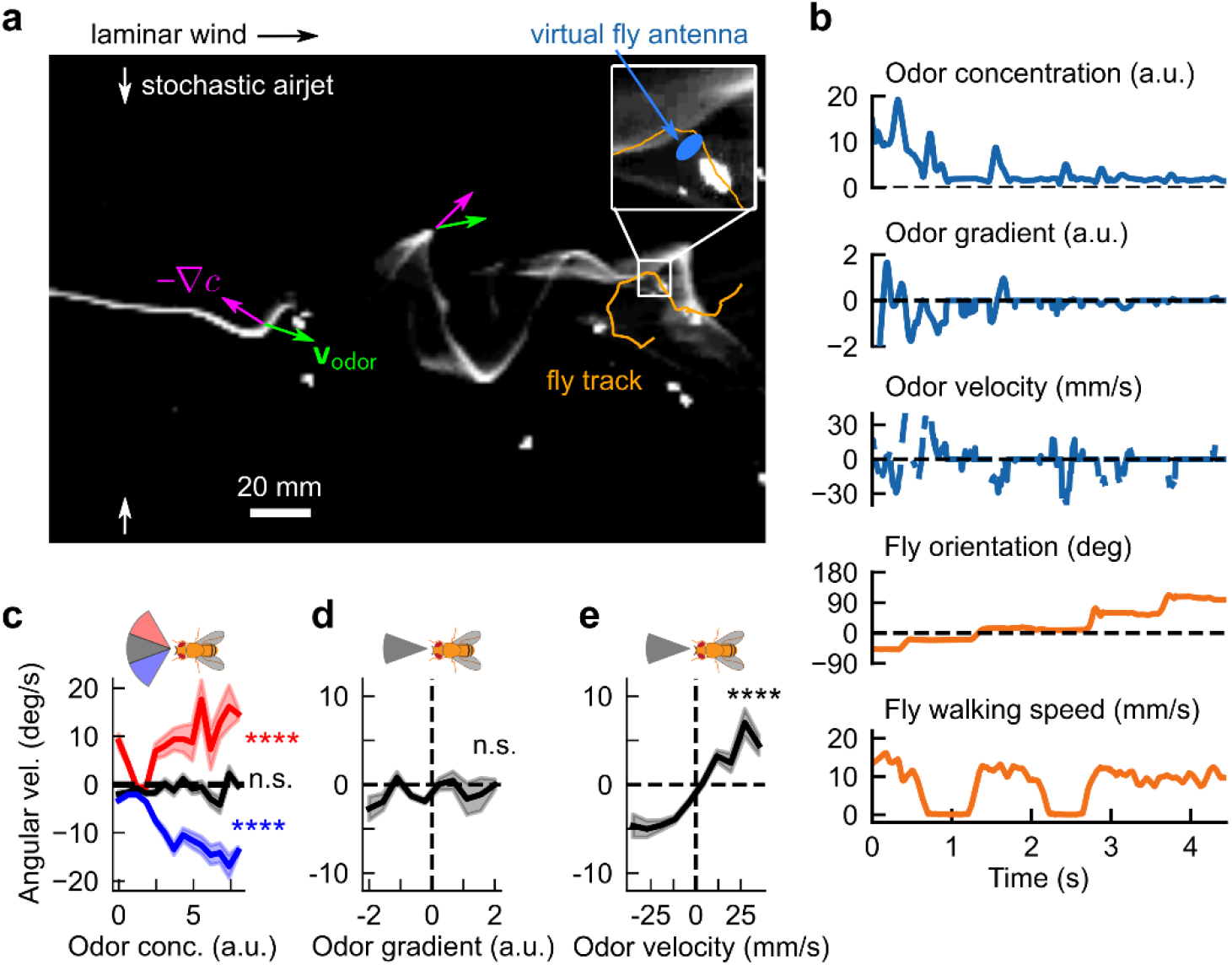
*Drosophila* turning behaviors are correlated with odor direction in a spatiotemporally complex odor plume. **a**, Snapshot of walking flies navigating a spatiotemporally complex odor plume generated by stochastically perturbing an odor ribbon in laminar flow with lateral airjets. Odor gradients (magenta arrows) and odor direction (green arrows) do not necessarily align, and can point in random directions relative to the odor source. Blue oval: virtual fly antennae region used to estimate perceived signal quantities during navigation. **b**, Example time trace of perceived signal-derived quantities (blue) and fly behaviors (orange) for track shown in **a**. Odor direction was computed by cross-correlating the signal in the virtual antenna over successive frames, and determining the spatial shift giving maximal correlation, while odor gradient was computed by linearly regressing the odor concentration against position along the major axis of the virtual antenna. **c**, Fly angular velocity as a function of odor concentration, for flies oriented in a 40° upwind sector (black), or in a 40° sector centered 20° clockwise (red) or counterclockwise (blue) from the upwind direction. Positive values indicate a counterclockwise turn. Correlations are significant for flies in the off-axis sectors (slopes = 0.037 ± 0.005, *n* = 174 tracks and −0.039 ± 0.003, *n* = 312 tracks for clockwise and counterclockwise sectors, respectively. *p* < 1*e*-6 (two-tailed t-test) for both sectors), but not those oriented directly upwind (slope = 0.005 ± 0.003, *p* > 0.05, *n* = 285 tracks). **d-e**, Fly angular velocity versus odor gradient and odor direction for flies oriented in a 40° sector upwind. Angular velocity is uncorrelated with odor gradient (mean slope = −0.005 ± 0.003, *p* > 0.05, two-tailed t-test, *n =* 284 tracks) but significantly correlated with odor direction (mean slope = 0.040 ± 0.003, *p* < 1*e*-6, two-tailed t-test, *n =* 282 tracks) in the virtual antenna.

Still, since odors are transported by the airflow, odor direction and wind motion are inherently correlated. To break this correlation, we turned to optogenetic stimulation of olfactory receptor neurons (ORNs) using the red-shifted channelrhodopsin Chrimson (Bell and Wilson, 2016; Klapoetke et al., 2014; Mafra-Neto and Cardé, 1994; Tao et al., 2020). We reasoned that not only would optogenetics allow us to adjust the airflow independently of the odor signal, it would also give us tight (< 300 µm) and fast (< 16 ms) control of the stimulus. We combined two experimental paradigms into a single optogenetic setup. The first is a large arena, high-throughput wind tunnel for walking fruit flies, also used to collect the data in Fig. 1 (Demir et al., 2020). The second is a method for patterned optogenetic stimulation using a light projector mounted above the arena (DeAngelis et al., 2020) (Fig. 2a). Our setup can deliver spatially complex light patterns throughout the arena, and individual flies can be optogenetically stimulated with sub-mm resolution. Due to Chrimson’s high sensitivity (Klapoetke et al., 2014), the relatively low light intensity of the projector (4.25 μW/mm^2^) over the large 27×17 cm^2^ arena was sufficient to stimulate a sustained firing response in ORNs, as verified with electrophysiology (Supplementary Fig. 2a). As a proof-of-concept, we projected fictive “odor ribbons” onto the arena while flowing laminar wind (Supplementary Fig. 2b), and recorded flies in which the olfactory co-receptor *Orco* drove the expression of Chrimson. Though flies are only weakly responsive to red light, we used blind flies throughout to remove any visual effects. Previous studies have shown that optogenetic stimulation of *Orco*-expressing neurons acts as an attractive fictive odor signal (Bell and Wilson, 2016; Tao et al., 2020). Indeed, flies turned and followed the fictive ribbons upwind, mirroring fly responses to streaming ribbons of attractive odors such as ethyl acetate and apple cider vinegar (Demir et al., 2020) (Supplementary Fig. 2b). By aligning the coordinate systems of the camera and projector, we can track flies’ behaviors simultaneously with their perceived fictive odor signal, giving us spatiotemporally precise measurements of fictive odor stimuli (Methods, Supplementary Fig. 2c).

**Figure 2.**
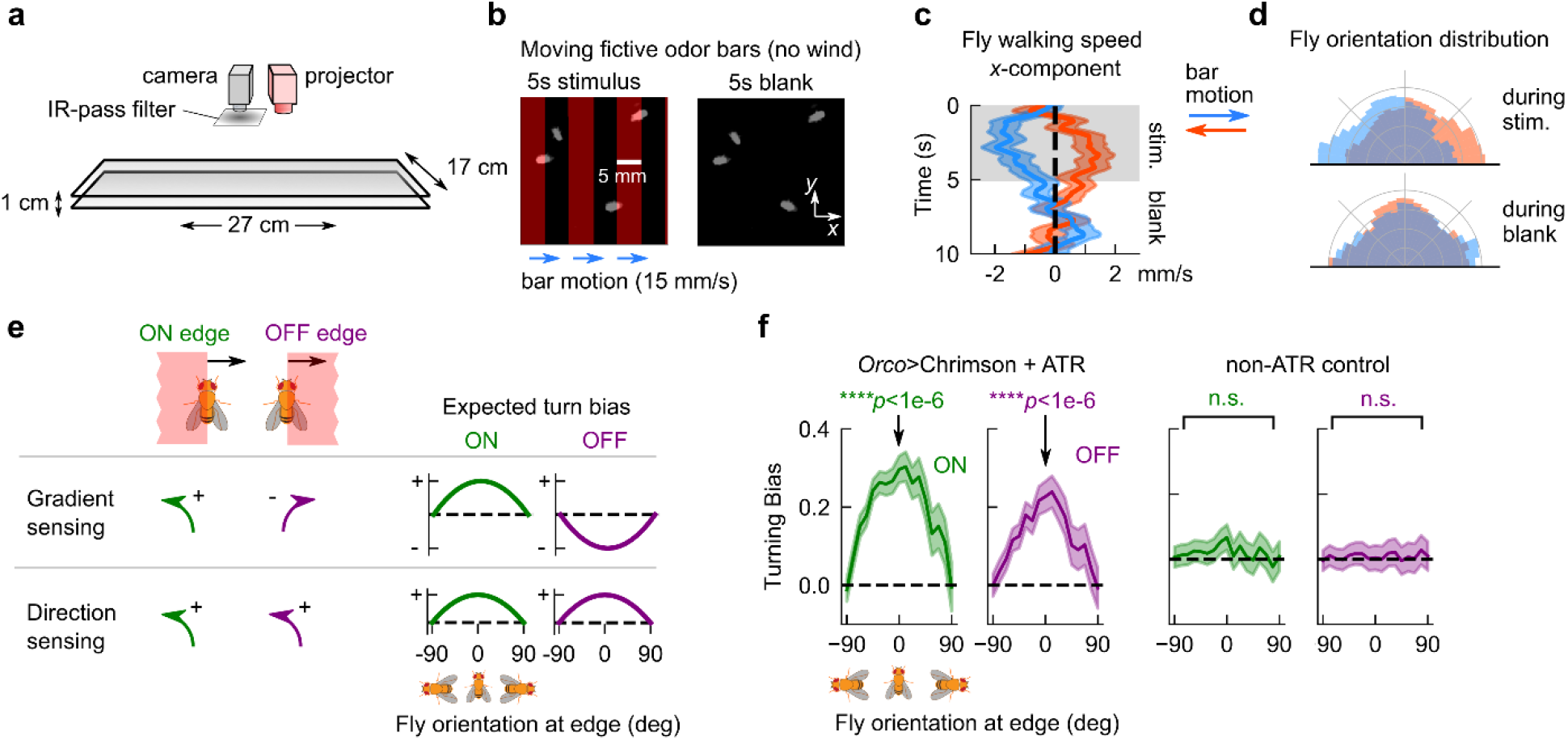
Turning responses are consistent with direction sensing, not gradient sensing. **a**, Schematic of fly walking assay. Flies with Chrimson expressed in ORNs receive optogenetic stimulation from a video projector mounted above arena, which displays fictive odor stimuli throughout arena with high spatial (< 300 um) and temporal (< 6 ms) precision. **b**, Fictive odor bars moving at 15 mm/s are presented in 5s blocks, interleaved with a 5s blank period. Differences in fly orientation or velocity for rightward (along +*x*) versus leftward (along -*x*) bar motion would indicate that flies can sense odor direction without mechanical cues from the wind. **c**, Component of fly walking velocity along +*x* direction during the 5s stimulus (shaded grey) and blank periods, for rightward (blue; *n* = 407 tracks) and leftward (orange, *n* = 455 tracks) moving bars, for *Orco*>Chrimson flies. Shaded error bars: SEM. **d**, Distribution of fly orientations during the 5s stimulus period (top) and 5s blank period (bottom), for rightward (blue) and leftward (orange) bar motion. Orientations are symmetrized over the *x*-axis. The differential effects in **c** and **d** disappeared for the same genotype with 1 antenna ablated (Supplementary Fig. 3a-b), but were maintained for flies with Chrimson expressed only in ORNs that express Or42b (Supplementary Fig. 3c-d). **e**, Direction sensing can be differentiated from gradient sensing by measuring turning responses as a function of fly orientation at both edges of wide, moving fictive odor bars: the ON edge (when the fictive odor passes onto the fly) and the OFF edge (when it leaves it). **f**, Fly turning bias versus orientation at ON (green) and OFF (purple) edge, for *Orco*>Chrimson flies that are optogenetically active (left 2 plots) and optogenetically inactive (i.e. not fed ATR; right 2 plots). Bars move at either 10 or 15 mm/s (data is pooled); turning bias is quantified as the sign of the change in orientation over the window from 150 ms to 300 ms after the bar onset, where +1 is counterclockwise and -1 is clockwise. Each point covers a span of ±45°; thus, distinct points contain overlapping data. Error bars: SEM. Turning bias for optogenetically active flies oriented perpendicular to the bar motion (*θ* = 0) are significantly distinct from zero for both ON and OFF edges (*p* < 1e-6 for both edges, chi-squared test; *n* = 2398 tracks), but not for optogenetically inactive flies (*p* > 0.05 for both edges; *n* = 3622 tracks).

Next, we presented a simple stimulus consisting of traveling fictive odors bars in the absence of wind. Flies oriented perpendicular to the bar motion receive differential stimulation across their antennae when the edges of each bar pass across them. If flies responded selectively to the direction of fictive odor motion, we would expect opposing behaviors for bars traveling rightward versus leftward. We thus presented 5mm-wide bars traveling 15 mm/s either left or right, in 5s-long blocks followed by a 5s-long block of no stimulus (Fig. 2b). Right-moving bars elicited a net displacement of fly position to the left, and vice versa (Fig. 2c). Further, flies oriented against the direction of motion during the 5s stimulus block, but exhibited no asymmetry during the 5s blank (Fig. 2d). Notably, both of these behaviors were absent in *Orc*o>Chrimson flies with one antenna ablated (Supplementary Fig. 3a-b), but were preserved when Chrimson was expressed only in ORNs expressing the receptor Or42b (Supplementary Fig. 3c-d), which is known to drive olfactory attraction to vinegar (Semmelhack and Wang, 2009). These experiments suggested that flies’ olfactory responses were direction selective, and that direction selectivity is enabled by bilateral sensing from the two antennae. The key indicator of direction selectivity was counterturning against bar motion – a reasonable response for locating an odor source emitting propagating odor signals.

### Direction selective responses to ON and OFF edges are computed with a timescale of tens of milliseconds

Since insects and vertebrates both detect spatial gradients of odor concentration and use them to navigate (Catania, 2013; Duistermars et al., 2009; Gardiner and Atema, 2010; Rajan et al., 2006; Wu et al., 2020), we wondered if gradient sensing could explain the directional biases we observed. We repeated the experiments above with wider (30-45 mm) bars, which allowed us to quantify responses to each edge individually – the ON edge, when the fictive odor first passes over the fly, and the OFF edge, when fictive odor leaves the fly (Fig. 2e). Responses to these stimuli would clearly distinguish direction selectivity from gradient sensing, since gradient sensing would result in opposing behaviors at the ON and OFF edges while direction sensing responses would be the same (Fig. 2e). We calculated fly turning bias, defined as the sign of the cumulative change in orientation between 150 and 300 ms after the edge hit, as a function of the fly’s orientation relative to the moving edge. For both ON and OFF edges, these plots had strong positive peaks for fly’s oriented parallel to the edge, indicating that flies are responding to the odor direction, not the spatial gradient (Fig. 2f). Meanwhile, the responses were flat for control flies (Fig. 2f). Repeating this for various bar speeds |***v***_bar_| showed strong direction selectivity for bars at 10 and 15 mm/s, and a suppression for lower speeds down to 1 mm/s (Supplementary Fig. 4). For slower speeds — 1 and 5 mm/s — the ON response was still significant, while the OFF response was absent, which could result from gradient sensing in nearly static odor environments. Finally, directional turning responses were essentially absent in two negative controls – flies in which Chrimson is not activated, or those with 1 antenna ablated (Supplementary Fig. 4).

### Turning responses to odor motion and wind motion are summed

Insects universally bias their heading upwind in the presence of odor (Alvarez-Salvado et al., 2018; Baker et al., 2018; Budick and Dickinson, 2006; Demir et al., 2020; Kanzaki et al., 1992; Kennedy and Marsh, 1974; Mafra-Neto and Cardé, 1994; Vickers and Baker, 1994), but the role of odor direction in this upwind response is unknown. Our patterned optogenetic setup allowed us to investigate this by independently controlling the wind and odor direction, which is otherwise impossible in natural environments. Above, we quantified turning bias in response to odor motion, but without wind (Fig. 2). We reasoned that in the presence of both wind and odor motion, fly responses would reflect some sort of summation of these responses in isolation, so we now presented fictive odors in wind, but without the motion of odor. To remove odor motion, we flowed laminar wind and flashed the entire arena for 2.5 seconds, followed by 2.5 seconds of no stimulus (Fig. 3a). This stimulates both antennae simultaneously, removing bilateral information — an artificial stimulus that is difficult to deliver with natural odors. In this situation, flies bias their heading upwind (against the wind) at the onset of the flash (Fig. 3b; left plot), reminiscent of their tendency to turn “against” the odor motion in the absence of wind (Fig. 2f). The similarity of turning responses to wind and odor motion separately is illustrated by fitting the turning bias versus orientation plots to a sinusoid (Fig. 3b; dashed lines). In both cases, the plots are well fit by *A*cos *θ*, where *A*_wind_ = -0.40 and *A*_odor_ = -0.30.

**Figure 3.**
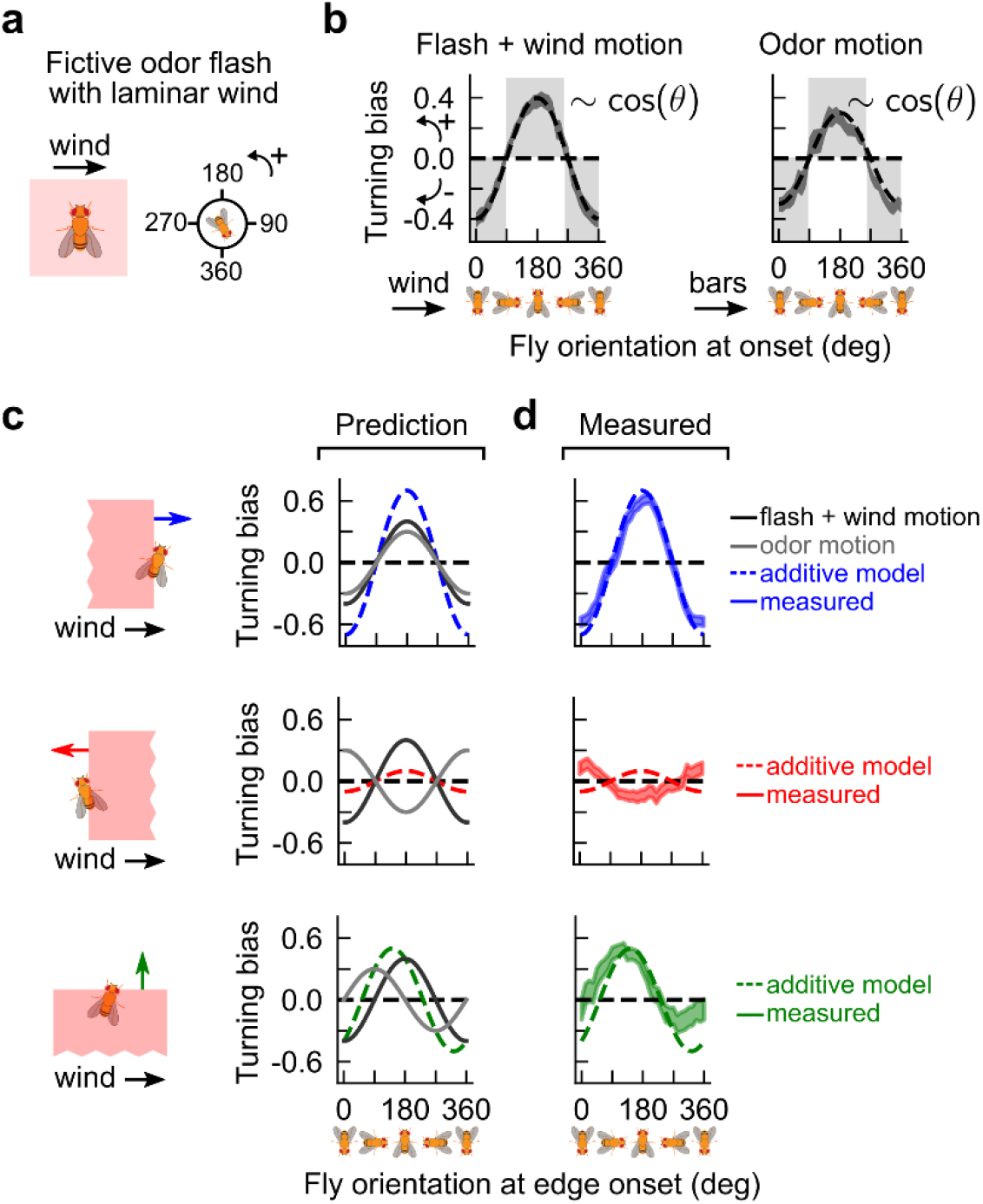
Turning responses to odor motion and wind motion are summed. **a**, Flashing the whole arena stimulates both antennae simultaneously, thus removing bilateral information that could enable direction selectivity. Laminar wind is introduced at 150 mm/s. **b**, Fly turning bias as a function of fly orientation, defined as in Fig. 2, for fictive bilateral odor flashes in the presence of wind (left) and moving fictive odor bars without wind (right). The latter plot is the same data as in Fig. 2f. Axes for the two plots are defined such that 90^0^ points in the direction of the wind or the direction of the bars, respectively. Grey shades: values for which fly turns counter to the wind direction or bar direction; all measured values lie in this range. Both plots can be well approximated by −0.4cos *θ* and −0.3cos *θ*, respectively. **c**, By row: expected turning bias versus orientation (dashed curve) for bars oriented parallel, antiparallel, or perpendicular to the wind, assuming that turning bias is the sum of the fitted cosines from **b**, which are reproduced in black and grey, respectively. Note that in the 2^nd^ and 3^rd^ row, the grey curve has a phase shift depending on the bar direction relative to the wind. **d**, Solid curves: measured data. Bars move at 15 mm/s. Dashed curves: expected responses from **c**. Shaded regions: 1 standard error. *n* = 2586, 2535, 2467, 1614 tracks for flash, and bars parallel, antiparallel, and perpendicular to the wind, respectively. Responses to OFF edges were very weak, suggesting other nonlinear interactions between the loss of odor and the wind (Supplementary Fig. 5).

These simple functional forms encouraged us to consider a simple hypothesis for how flies respond to fictive odor edges moving at a given angle relative to the wind. We hypothesized that the response to the combined signal is a sum of the bar motion and odor motion responses. This hypothesis predicts that when the odor and wind direction are aligned, the peak response should increase in magnitude and remain centered at 0° and 180° (Fig. 3c; first row). If odor and wind motion oppose each other, these peaks should nearly cancel (Fig. 3c; middle row). Finally, in the interesting case of wind and odor directions perpendicular to each other, the peaks should shift leftward to ∼145° and ∼325° (Fig. 3c; bottom row). To test these predictions, we presented fictive odor bars either parallel, antiparallel, or perpendicular to 150 mm/s laminar wind. When the wind and odor were aligned, the turning bias at ON edges was nearly perfectly fit by the additive prediction (Fig. 3d). The antiparallel motion of bars and odors was also fit well – extrema remained at 0° and 180°, though the cancellation overshot slightly. Notably, the response to perpendicularly oriented wind and odor reproduced the shift of the response curve peak from ∼180° to 145°, and nearly reproduced the shift of the minimum from ∼360° to ∼325°. These results suggest that odor direction selective responses integrate with directional information from the wind in a largely, but not entirely, additive fashion. Moreover, universally observed upwind turning responses are more than naive mechanosensory reactions triggered by the presence of odor – they can be enhanced and even cancelled by directional information from the odor itself.

### Flies use spatiotemporal correlations in odor intensity to detect odor direction

We next tested the extent to which our observations were consistent with elementary motion detection algorithms, by first analyzing our data for moving bars in the absence of wind (Fig. 2). Odor motion creates a difference in latency Δ*T* between the stimulation of the two spatially separated antennae, the sign and magnitude of which determines the output of direction-selective models such as the classical Hassenstein-Reichardt correlator (HRC) (Hassenstein and Reichardt, 1956). In our assay, Δ*T* can be inferred from the velocity of the bars relative to the flies using simple geometric considerations (Supplementary Fig. 6; Methods). This allows us to express turning bias as a function of Δ*T*, thereby directly testing the predictions of an HRC model. In a rightward-selective HRC (Fig. 4a), a signal from the left antenna is multiplied with the delayed signal from the right antenna, where the delay is implemented as an exponential filter *e*^−*t*/*τ*^. Subtracting this from a similar computation with the antennae switched gives the detector output *r*(*t*). We modeled the turning bias as the time integral of *r*(*t*), for which the HRC predicts a turning bias proportional to 1 − *e*^−Δ*T*/*τ*^ for rightward moving edges. Thus, plotting the turning bias against Δ*T* would allow us to extract the filter time constant *τ*, revealing the timescale of olfactory motion detection. Pooling the data from both ON and OFF edges, we found that the prediction was fit well, with filter timescales in the range *τ* = 25 ± 12 ms (Fig. 4b). Though this estimate is approximate and limited by the temporal and spatial resolution of the projector, it is notable that the timescale is comparable to the timescales of visual motion detection in *Drosophila* vision (Salazar-Gatzimas et al., 2016).

**Figure 4.**
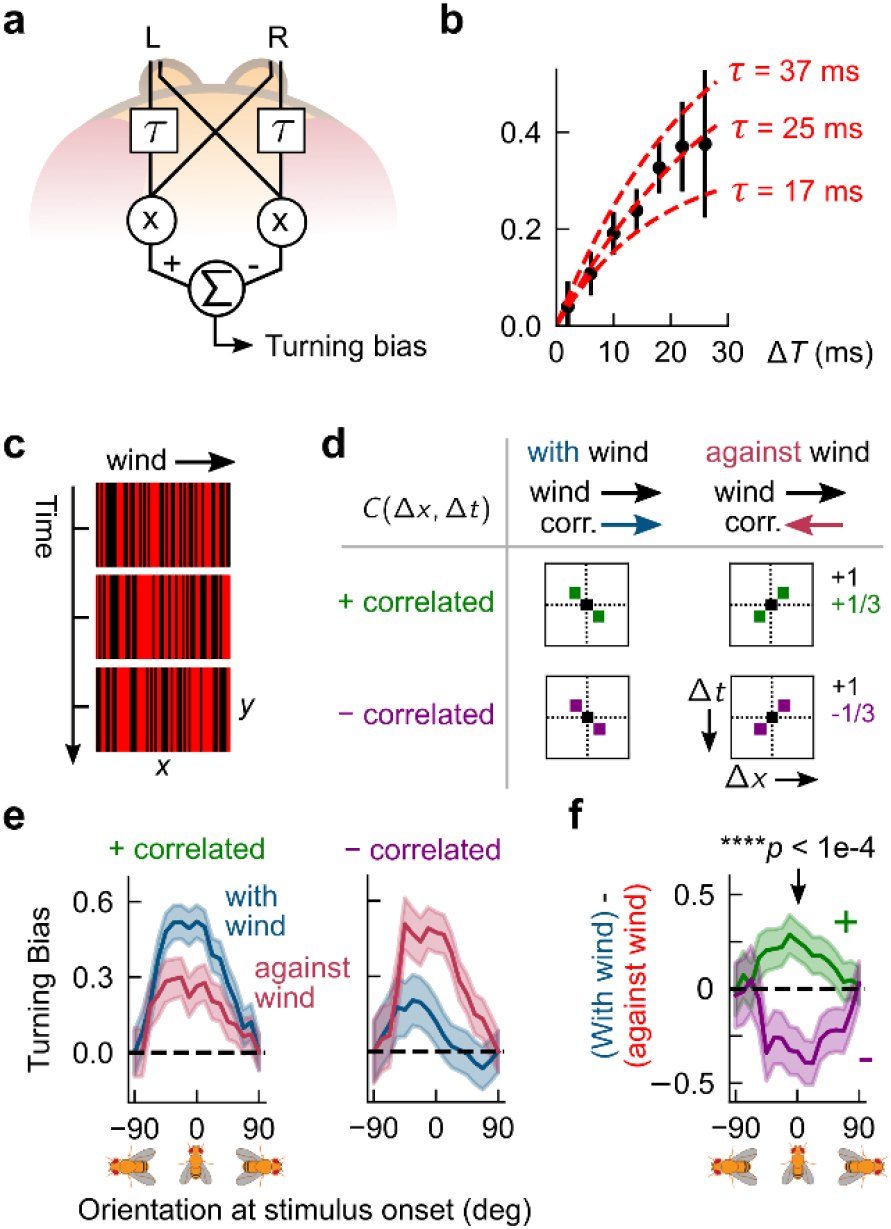
Olfactory direction sensing obeys a correlation-based algorithm. **a**, Schematic of hypothesized Hassenstein-Reichardt correlator (HRC) model in the olfactory circuit. Signal from one antenna projects to both brain hemispheres, but with distinct temporal transformations; we implement this by filtering one arm with *e*^−*t*/ *τ*^. Fly’s turning bias is modeled as the time integral of the correlator output (Methods). **b**, Black dots: measured turning bias versus Δ*T*, for all times fly crosses a fictive odor edge. Each datapoint spans ±4 ms. The HRC model predicts that turning bias is proportional to 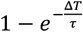, which can be used to extract the delay timescale *τ*. Middle red line: fit of HRC to mean of turning bias; upper/lower lines: fit to mean ± 1 SEM of the turning bias. Estimated correlator timescale *τ* lies in a range of tens of milliseconds. **c**, Correlated noise stimuli consist of 1-pixel-wide fictive odor bars perpendicular to 150 mm/s laminar flow. In one frame, each bar is independently bright or dark with equal probability (3 subsequent frames are shown). However, stimuli are correlated in time, so the bar pattern in the next frame depends on the pattern in the current frame. In this illustration, bars are positively correlated along +*x*, so a bright bar at a given *x* in one frame is likely to be proceeded by a bright bar one *x*-pixel to its right in the next frame. Visually, this would look like a rightward moving pattern. **d**, There are 4 types of stimuli, depending on the correlation direction (along +*x*, i.e. with-wind, or along – *x*, i.e. against the wind) and the correlation parity (+ or -). Each type of stimulus is characterized by the correlation matrix *C*(Δ*x*, Δ*t*) between two bars separated spatiotemporally by Δ*x* pixels and Δ*t* frames. Since our stimuli are generated by summing and binarizing Gaussian variables, nonzero correlations are not absolute, but rather have magnitude 1/3. For example, for positively correlated with-wind stimuli (top left plot), *C*(1, 1) = *C*(−1, −1) = 1/ 3, and the remaining correlations are zero, while for negatively correlated with-wind stimuli (bottom left plot), *C*(1, 1) = *C*(−1, −1) = −1/ 3. **e**, Turning bias versus fly orientation for positively correlated (left) and negatively correlated (right) stimuli. Stimuli are presented in 4s blocks, interleaved with 4s of no stimulus; wind flows throughout. Turning biases are defined as the sign of the change in orientation over 300 ms from the onset of the 4s stimulus block. *n* = 489, 496 for positively correlated with and against-wind, and 338, 335 for negatively correlated wind and against-wind, respectively. **f**, Difference *D* between with-wind and against-wind responses from **c**, for positively (green) and negatively (purple) correlated stimuli. The value of *D* for positive and negative correlations differed significantly for flies oriented perpendicular to the bar motion (*θ* = 0), (*p* < 1e-4, chi-squared test).

Elementary motion detection algorithms respond fundamentally to correlations in the signal over space and time. To better compare against the predictions of the HRC, we moved beyond ON and OFF odor edges and turned to *correlated noise* stimuli, which have been used to characterize direction selective computations in fly vision (Salazar-Gatzimas et al., 2016). A snapshot of a correlated noise stimuli is a pattern of 1-pixel wide bars, each of which is either bright or dark (Fig. 4c). The pattern updates in time in such a way that it contains well-defined positive or negative correlations between adjacent pixels. Intuitively, a positive correlation in the +*x* direction means that bright bar at a given *x* is likely to be proceeded, in the subsequent frame, by a bright bar 1 pixel to its right; visually, this would appear to be a rightward moving pattern. To enhance the effects, we simultaneously flowed laminar wind as in the experiments in Fig. 3. Thus, there were four types of correlated noise stimuli, corresponding to the possible combinations of correlation direction (with or against wind) and polarity (negative or positive), each of which is uniquely defined by its correlation matrix *C*(Δ*x*, Δ*t*) (Fig. 4d).

In this experiment, turning responses to positively-correlated noise stimuli mimicked those to moving bars: upwind turning was suppressed when the correlation direction opposed the wind (Fig. 4e; first plot). Importantly, spatial gradients in these stimuli quickly average to zero, so only a computation sensitive to spatiotemporal correlations — and not gradients — could account for behavioral suppression when the correlation direction and wind were misaligned. Repeating for negative correlations, we found that upwind turning was suppressed when the correlation and wind were *aligned* (Fig. 4e; second plot). Notably, this response is also consistent with a correlation-based algorithm, which predicts a reversal of behavior when the correlation polarity flips sign (Salazar-Gatzimas et al., 2016). In fact, this “*reverse phi*” phenomenon is actually an illusion – a byproduct of a pairwise correlator algorithm – that has been observed in visual responses of several species (Clark et al., 2011; Livingstone et al., 2001; Orger et al., 2000; Salazar-Gatzimas et al., 2018; Tuthill et al., 2011), including humans (Anstis and Rogers, 1975). Subtracting the with-wind and against-wind responses for each polarity indicated clearly that the reverse phi prediction was satisfied (Fig. 4f).

We corroborated our results using *gliders*, another class of correlated stimuli (Clark et al., 2014; Hu and Victor, 2010). Visually, a glider is a random pattern of light and dark bars moving in one direction (Supplementary Fig. 7a). Unlike correlated noise, the bars are correlated not only with a neighboring bar in the subsequent frame, but also with more distant bars at later times. However, unlike the weaker 1/3 correlations for correlated noise, the correlations in glider stimuli are perfect (Supplementary Fig. 7b), so we expected similar trends as before, but with larger effect sizes. For positively correlated gliders, we found similar trends as with correlated noise, but much larger separations between the with-wind and against-wind responses (Supplementary Fig. 7c). We were also able to explore a range of correlation times by adjusting the frame update times. For update times in the range of 17-30 ms, we find direction selective responses, while for shorter update times (11 ms), direction selectivity disappeared (Supplementary Fig. 7d). Interestingly, the maximum separation of with-wind and against-wind responses was with a frame update of 17-22 ms, consistent with the estimate of the HRC filter constant using moving bars (Fig. 4b).

For flies to sense these correlations in our assay, their antennae must be optogenetically stimulated by distinct pixels. We satisfied this requirement by mounting the projector such that the *x*-pixel width (∼290 µm) approximated the *D. melanogaster* antennal separation (Supplementary Fig. 7e) (Miller and Carlson, 2010). Consistent with this, effects must also reduce for bars that are wider than the antennal separation. Indeed, repeating the experiments with double the bar width, we found no significant differences between with-wind and against-wind responses (Supplementary Fig. 7f). Together, these results suggest that *Drosophila* olfactory direction sensing obeys a correlation-based algorithm.

### Odor direction encodes crosswind position and aids navigation in complex plumes

Animals could use measurements of odor direction to help them navigate complex plumes, provided this information complements other directional cues such as gradients or wind. To quantify the distribution of odor signal directions in a naturalistic plume, we ran numerical simulations of an environment replicating the plume from Fig. 1. These simulations provide not only a more finely resolved concentration field, but also the airflow velocity field (Fig. 5a), which is experimentally inaccessible. We first compared, for a few fixed points in the plume, the odor velocity **v**_odor_ and the airflow **v**_wind_ at a single time. Both **v**_odor_ and **v**_wind_ had *x*-components comparable to the mean flow speed 150 mm/s. However, **v**_odor_ also had large crosswind components **v**_*y*,odor_ pointing outward from the plume centerline, which were noticeably absent from **v**_wind_ (Fig. 5b; left). Averaging over all detectable odor filaments in the 120s simulation revealed a similar trend: away from the plume centerline, the distribution of **v**_odor_ spanned a tight angular range, pointing consistently outward in the crosswind direction (Fig. 5b; middle column). Meanwhile, ***v***_wind_ was distributed largely downwind, with much smaller outward angles (Fig. 5b; right column). To visualize the “flow” of odor motion, we calculated the time-average of ⟨**v**_odor_⟩ at all locations in the plume. We compared this to the time-average of the wind vector conditional on the presence of odor, ⟨**v**_wind|odor_⟩. We used the latter rather than the unconditional wind velocity, ⟨**v**_wind_⟩, since for an ideal point source of odor within homogeneous turbulence, the latter does not encode the lateral location of the source. Throughout the plume, ⟨**v**_odor_⟩ flowed strongly outward from the plume center, while ⟨**v**_wind|odor_⟩ was directed essentially downwind (Fig. 5c).

**Figure 5.**
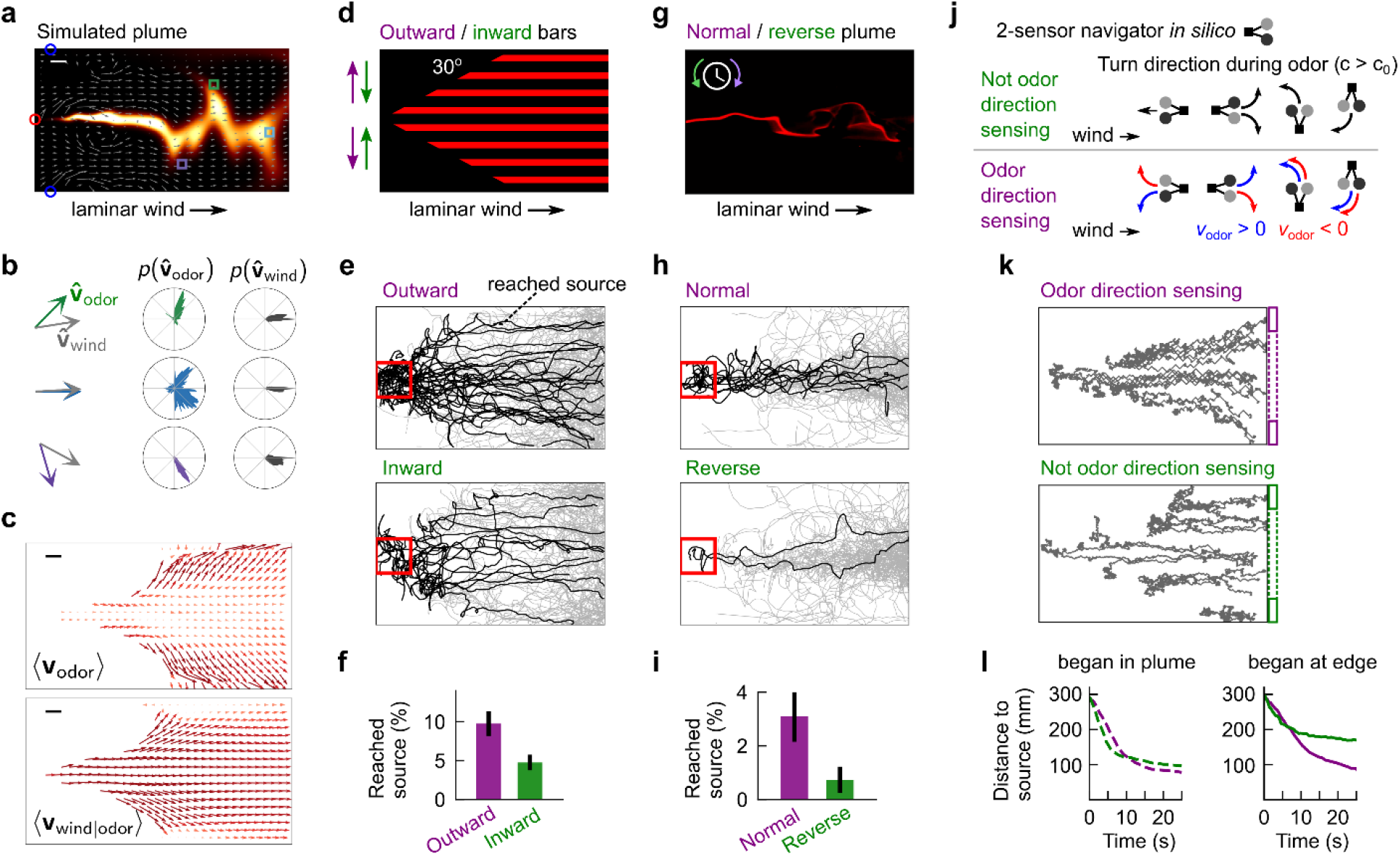
Odor direction detection enhances natural plume navigation. **a**, Snapshot of direct numerical simulation of complex odor plume from Fig. 1. Grey vector field: airflow at snapshot instant; white scale bar: 20 mm. **b**, (left column) Odor velocity vector at corresponding boxed locations in **a** along with airflow direction vector at same position. (middle column) Histogram of odor velocity at all times in simulation, at corresponding positions in **a**. (right column) Same for wind. **c**, (top) Odor velocity vector field, averaged over entire simulation. (bottom) Vector field of wind velocity, for times at which odor concentration is detectable, averaged over entire simulation. Vectors are colored by magnitude from low (yellow) to high (maroon). **d**, Illustration of fictive odor landscape in which bars move laterally outward or inward from center of the arena. Bars are restricted to a conical region approximating the envelope of a complex plume emanating from a source. Laminar wind flows at 150 mm/s. Experiments used 2 mm wide bars moving at 15 mm/s and spaced by either 5, 10, or 15 mm (data is pooled); these gave fictive odor hit frequencies in the range ∼1-2 Hz, similar to the measured plume. **e**, Measured tracks for flies beginning in the rear 50 mm of the arena, navigating the plume depicted in **d**, for outward (top) and inward (bottom) moving bars. Black tracks: fly tracks that reached a 40 mm box around the fictive plume source. *n* = 312, 457 tracks for outward and inward bars. For visual comparison, the same number of tracks (312) are shown in both plots. **f**, Percentage of tracks beginning in rear 50 mm that reached the source (red box in **e**); means are 9.8% and 4.8% for the outward and inward plumes, respectively. SEMs determined by bootstrapping over individual trajectories; differences are significant (*p* < 0.01, two-tailed t-test). **g**, Snapshot of recorded plume from Fig. 1, optogenetically projected into the arena with normal playback or reverse playback. Reversing the playback preserves the spatial location of odor hits and other temporal features, but reverses the local odor direction. **h**, Measured tracks for flies navigating the complex plume depicted in **g**, when the video is played normally (top) or in reverse (bottom). Only considered are tracks beginning in the rear 50 mm of the arena and within 30 mm laterally from the plume centerline; further from the centerline, there is no detectable stimulus. *n* = 295 and 277 tracks for normal and reverse playback, respectively. **i**, Percentage of tracks that reached the source; means are 3.0% and 0.7% for forward and reverse playback, respectively; differences are significant (*p* < 0.05, two-tailed t-test). **j**, 2-sensor robot navigator *in silico*. Agents are always oriented at 0°, 90°, 180° or 270°, and at each timestep turn 90° either left or right and move forward one step. Agents are either direction sensing (DS+) or not direction sensing (DS-) When odor concentration *c* exceeds some threshold *c*_0_, DS-agents turn upwind. DS+ agents, for *c* > *c*_0_, turn against the odor direction when oriented upwind or downwind; crosswind agents always turn upwind. DS+ agents infer odor direction using a simple spacetime correlation between their 2 sensors (Methods). **k**, Example trajectories of robots navigating plume in **a**, when they are initialized in the back of the arena. **l**, Distance to source over time, for those with (purple) and without (green) odor direction sensing ability, for robots initialized near the plume centerline (<120 mm from axis; left plot) or near the plume edges (right plot). DS+ agents make significantly quicker progress when initialized near the plume edges.

This analysis suggests that in naturalistic odor plumes emanating from a point source, odor direction is a strong indicator of the direction towards the centerline of the plume. This directional cue is not necessarily reflected in the local wind, nor in the local gradients, though we did find that odor gradients have a similar crosswind structure closer to the source, where the plume is less intermittent (Supplementary Fig. 8a). Of course, to be useful for navigation, odor direction must be resolvable on realistic timescales. By calculating the running average of the odor direction at a fixed location, we found that in most of the plume extent, only several hundred milliseconds were necessary to resolve the lateral components (Supplementary Fig. 8b-c). Since odor bursts occurred at ∼1-5 Hz in this particular plume, a navigator could estimate the direction of odor motion orthogonal to the mean flow after only a few odor hits.

To investigate how *Drosophila* use odor motion during a navigation task, we designed a fictive odor plume whose boundaries were subtended by a cone — as if emanating from a source — and within which thin bars moved laterally outward from or inward toward the centerline, while laminar wind flowed along the cone axis (Fig. 5d). We reasoned that inward moving bars, which are reversed from their natural flow, would degrade localization to the odor “source,” i.e. the tip of the cone. For both bar directions, flies stayed within the conical fictive odor region, but were significantly more likely to reach the upwind source region when the bars moved naturally outward (9.8% versus 4.8% reached the source for outward versus inward bars, respectively, *p* < 0.01, two-tailed t-test) (Fig. 5e-f). Notably, the fictive odor signals in these two paradigms do not differ by location, frequency, duration, or spatial gradient — differences in performance (Fig. 5f) can only be explained by odor direction alone. We then tried the more realistic case of projecting a video of a recorded plume (Fig. 1a) onto the arena (Fig. 5b), playing the video not only normally, but also in reverse. As in the previous paradigm, reverse playback reverses odor direction without perturbing any other spatial or temporal information measured at each point. Remarkably, the likelihood to reach the odor source significantly degraded when the plume was played in reverse (3.0% versus 0.7%; *p* < 0.05, two-tailed t-test) (Fig. 5h-i). Together, these results indicate that the odor motion provides a directional cue complementary to odor gradients and wind motion, and strongly enhances navigation in complex odor plumes, even when all other aspects of the odor signal remain unchanged.

Finally, with an eye toward practical applications, we used *in silico* experiments to explore the impact of odor motion sensing for robots obeying a simplified navigation algorithm. Virtual agents detected odor signals using two spatially separated olfactory “sensors,” from which they inferred odor direction *v*_odor_ = ±1 using a rudimentary HRC-like computation (details in Methods). We simulated two types of agents, with and without odor direction sensing (DS+ and DS-agents, respectively). Agents were always oriented in one of the 4 cardinal directions; at each frame, they turned 90° either left or right and moved forward one step. For undetectable odor concentrations (odor concentrations *c* less than some threshold *c*_0_), turns were randomly left or right with equal probability. For DS-agents, navigation followed a simple odor-gated anemotaxis strategy, in which agents moved upwind in the presence of odor. Specifically, for *c* > *c*_0_, crosswind agents turned upwind, upwind agents maintained their heading, and downwind agents turned randomly left or right (Fig. 5j; first row). DS+ agents, on the other hand, obeyed a combination of odor-gated anemotaxis and odor-direction-biased taxis. Specifically, odor-elicited turns were shaped by odor direction whenever the wind provided no bias (Fig. 5j; second row). Thus, for *c* > *c*_0_, crosswind agents still turned upwind, but those facing up- or downwind turned “against” the odor motion (left or right turns for *v*_odor_ = 1 or *v*_odor_ = −1, respectively), provided the odor motion was above a detectable threshold.

Putting these agents in the simulated plume (Fig. 5a), we found that both DS+ and DS-agents starting in the back of the arena could eventually find their way to the odor source (Fig. 5k). In particular, both fared well when initialized near the plume axis – in fact, DS-agents reached the source slightly more efficiently, unhindered by suboptimal crosswind moves when already facing upwind (Fig. 5l; dashed line). However, if initialized closer to the plume edges, DS-agents’ progress quickly deteriorated once they surpassed the conical extent of the plume (Fig. 5k-l). Meanwhile, DS+ agents were aided by lateral motion toward the plume axis (Fig. 5k), leading to significantly more sustained progress toward the source (Fig. 5l). This indicated that the clearest benefit of odor direction sensing was an increase in navigation reliability for sub-optimal starting positions. Thus, even a simplistic implementation of odor motion sensing can enhance the robustness of complex plume navigation, and could be incorporated straightforwardly to olfactory robots in a variety of existing schemes (Gumaste et al., 2020; Hengenius et al., 2021; Kowadlo and Russell, 2008; Liu et al., 2020; Riman et al., 2021).

## DISCUSSION

Olfactory navigation relies on integrating various sensory signals that contain information about the odor source. Which features exist, and how much information they carry, can vary considerably between plume structures (Boie et al., 2018; Jayaram et al., 2021; Rigolli et al., 2021). Gradient sensing can provide reliable directional information when navigating laboratory-controlled plumes, such as static ribbons (Duistermars et al., 2009), or very close to the source of natural plumes before odor patches have dispersed (Supplementary Fig. 8). Further away from the source however, turbulent air motion stretches and fragments odor regions as they are carried downstream, producing odor signals that are patchy and intermittent (Celani et al., 2014; Riffell et al., 2008), and which span many spatial scales – the so-called inertial convective range – from macroscopic eddies to molecular diffusion (Sreenivasan, 2019). In these regions, odor concentration gradients tend to point in random directions relative to the source, and so have limited value. Even in turbulent boundary layers, where concentrations are more regular (Connor et al., 2018), gradients can aid navigation, but require unnaturally amplifying the gradient to an extreme degree not consistent with data (Alvarez-Salvado et al., 2018).

Our work confronts some of the limitations of gradients by revealing an entirely distinct role for bilateral sensing: measuring odor direction by comparing concentrations in both space and time. This information stream is especially relevant to the statistical features present in the inertial convective range of turbulent plumes. Parallel to the plume axis, odor motion is mainly determined by, and redundant with, the average wind direction. But perpendicular to the plume axis, turbulence spreads odor packets by random continuous motions, with an effective diffusivity much larger than molecular diffusion (Pope, 2011; Taylor, 1922). What results is a flux of odor patches directed away from the plume centerline, providing a strong directional cue orthogonal – and thus complementary – to the mean wind. We corroborated this with theoretical analysis of a simple turbulent plume model (Methods), finding that the outward flow of odor motion we found in simulations (Fig. 5c) exists in turbulent odor plumes more generally (Supplementary Fig. 9a-b), and that lateral odor velocity components can be detected by computing local correlations between two nearby points (Supplementary Fig. 9c).

Insects universally bias their heading upwind when odors become longer, more intense, or more frequent (Alvarez-Salvado et al., 2018; Baker et al., 2018; Demir et al., 2020; Kanzaki et al., 1992; Kennedy and Marsh, 1974; Mafra-Neto and Cardé, 1994; van Breugel and Dickinson, 2014). This strategy fails at the plume edges, where insects then resort to local search or downwind or crosswind motion to re-enter the plume (Alvarez-Salvado et al., 2018; Budick and Dickinson, 2006; Mafra-Neto and Cardé, 1994). In this sense, the value of the lateral odor motion is evident, providing cues about which crosswind direction to take to reenter the plume. Our work does not explore odor direction sensing in the z-dimension – say, for flying insects. The role of odor direction sensing would likely be different, since odors traveling upward would not be sensed bilaterally unless the fly were flying with nonzero roll. In flight, directional cues from the optic flow also shape navigation (Budick et al., 2007). How odor direction contributes in this locomotor regime remains an avenue for future work.

Our setup allows us to test the predictions of the HRC using artificial correlation-type stimuli which would be prohibitive to reproduce with natural odors. In particular, we generated a *reverse phi* illusory percept for negative correlations, an signature of correlation-based algorithms observed in visual motion detection in flies (Clark et al., 2011; Eichner et al., 2011; Salazar-Gatzimas et al., 2018; Salazar-Gatzimas et al., 2016; Tuthill et al., 2011) and other species (Hassenstein and Reichardt, 1956; Livingstone et al., 2001; Orger et al., 2000), including humans (Anstis and Rogers, 1975). The HRC computes only second-order correlations – correlations between pairs of points in space and time – but, at least in vision, higher-order correlations can elicit direction-selective behaviors (Clark et al., 2014), and may improve motion detection by exploiting the statistics of natural scenes (Chen et al., 2019; Fitzgerald and Clark, 2015; Fitzgerald et al., 2011). Natural odor landscapes also exhibit universal highly-structured statistics (Celani et al., 2014) to which odor direction selective computations may likewise be tuned.

In mouse retina and fly vision, motion detection circuits have been characterized in detail and have many parallels (Borst and Helmstaedter, 2015; Clark and Demb, 2016), though much remains unknown. In both, visual motion is computed separately for ON and OFF edges (Euler et al., 2002; Famiglietti, 1983; Maisak et al., 2013), and it is likely that a similar split may exist in odor motion computations, given the difference in responses to ON and OFF edges in the presence of wind (Supplementary Fig. 5). In contrast to the canonical HRC architecture, three inputs feed into direction selective neurons in the fly visual circuit (Shinomiya et al., 2019; Takemura et al., 2017). This is unlikely to be the case in olfaction, if direction sensing is indeed enabled by bilateral segregation. Still, our results do not implicate any specific circuit architecture or mechanism. In fly vision, direction selective behaviors and signals are frequently well-described by a pairwise correlator model (Clark et al., 2011; Haag et al., 2004), while the underlying neural architectures and functional interactions remain incompletely understood and quite complex (Badwan et al., 2019; Gruntman et al., 2018, 2019; Haag et al., 2016; Salazar-Gatzimas et al., 2018; Shinomiya et al., 2019; Strother et al., 2017; Takemura et al., 2017; Wienecke et al., 2018). Ultimately, comparisons between odor and visual motion detection systems will reveal how circuits in these distinct modalities accomplish similar tasks.

Where could direction selectivity occur in the olfactory circuit? Most ORNs project to both antennal lobes, but ipsilateral and contralateral signals differ in magnitude and timing (Gaudry et al., 2013; Tobin et al., 2017), which could be amplified further downstream to enact bilateral computations. One potential region of interest is the third-order olfactory center, the lateral horn (LH), which mediates innate odor responses (Jefferis et al., 2007). Output neurons from the LH to the ventrolateral protocerebrum (VLP) have been shown to enhance existing bilateral differences through contralateral inhibition (Mohamed et al., 2019). Though this may be an isolated effect, the VLP region is highly suggestive: it lives in the ventral region of the LH, which receives inputs from wind-sensing wedge neurons – a potential integration center for bilateral odor information and wind (Dolan et al., 2019).

The lack of smooth concentration fields in naturalistic plumes has inspired a number of studies focusing on how animals use the temporal features of the odor signal, such as the frequency of encounters with odorized air packets. This reliance on timing is enabled by the remarkable degree of temporal precision in olfactory circuits (Ackels et al., 2021; Gorur-Shandilya et al., 2017; Martelli et al., 2013; Park et al., 2016; Shusterman et al., 2011). Here, we show that odor timing can be combined with spatially-resolved sensing to produce a complementary information stream, encoding directions that do not exist in the only other directional cue, the wind. Our work reveals a novel role for bilateral sensing in turbulent plume navigation, beyond measuring simple gradients.

## ACKNOWLEDGEMENTS

We thank Brian DeAngelis for helpful conversations and advice on the design of the optogenetic component to the walking assay, Omer Mano for help with projector troubleshooting, and Aarti Sehdev for help with behavioral experiments, fly rearing, and discussions. We also thank Viraaj Jayaram, John Carlson, Jamie Jeanne, and members of the Emonet Lab for helpful discussions and advice on the project. NK was supported by a postdoctoral fellowship through the Swartz Foundation for Theoretical Neuroscience, by postdoctoral fellowship NIH F32MH118700, and by postdoctoral fellowship NIH K99DC019397. MD and TE were partially supported by the Program in Physics, Engineering, and Biology at Yale and by NIH R01GM106189 and R01GM138533. BM and MR were supported by National Science Foundation grant IIS-1631864. DAC was supported by NIH R01EY026555 and NIH R01NS121773.

## CONTRIBUTIONS

NK, DC and TE designed the research. NK and MD built the assay with inputs from DC and TE. NK performed all experiments, data analysis, and agent-based simulations. MD performed the electrophysiology. BM and MR performed the numerical simulations for Fig 5. NK and TE performed the theoretical analysis of the turbulent plume. NK, DC and TE validated the data. NK, DC, and TE discussed the data analysis. NK, DC, and TE wrote the initial draft and all revisions. All authors approved the final manuscript.

## METHODS

### Fly strains and handling

Flies were reared at 25°C and 60% humidity on a 12 hour/12 hour light-dark cycle in plastic vials containing 10 mL standard glucose-cornmeal medium (i.e. 81.8% water, 0.6% agar, 5.3% cornmeal, 3.8% yeast, 7.6% glucose, 0.5% propionic acid, 0.1% methylparaben, and 0.3% ethanol. Media was supplied by Archon Scientific, NC). All flies used in behavioral experiments were females. Between 10 and 30 females were collected for starvation and placed in empty vials containing water-soaked cotton plugs at the bottom and top. All flies were 3–10 days old and 3 days starved when experiments were performed. Optogenetically active flies were fed 1 mM all trans-Retinal (ATR) (MilliporeSigma; previously Sigma Aldrich) dissolved in water. ATR was fed to flies 1 day prior to recording.

All flies used throughout the study had copy of the *GMR*-*hid* gene to make them blind. Optogenetic activation was achieved by expressing Chrimson (20X-UAS-CsChrimson) in *Orco*-expressing olfactory receptor neurons (Orco-GAL4) in almost all experiments. The one exception was the single-Or experiments (Supplementary Fig. 3c-d), which expressed Chrimson in only neurons expressing the olfactory receptor Or42b.

### Behavioral assay and optogenetic stimulation

The fly walking assay is identical to the one used in a previous study (Demir et al., 2020). All experiments were done in a behavioral room held at 21-23°C and 50% humidity. The walking arena is 270×170×10mm (see Fig. 2a), and consists of top and bottom glass surfaces and acrylic sidewalls. The upwind end is an array of plastic coffee straws, which laminarize the airflow (when wind is turned on); downwind end is a plastic mesh. For experiments with wind, dry air is passed through the straws at a flow rate giving a laminar flow at 150 mm/s within the arena. Flies are introduced by aspirating through a hole near the downwind plastic mesh. Flies were illuminated using 850 nm IR LED strips (Waveform Lighting) placed parallel to the acrylic sidewalls.

Experiments were recorded with a FLIR Grasshopper USB 3.0 camera with IR-pass filter at 60 Hz. Optogenetic stimuli were delivered using a LightCrafter 4500 digital light projector mounted 310 mm above the arena, illuminating an area larger than in the original method (DeAngelis et al., 2020). Only the red LED (central wavelength 627 nm) was used throughout this study. We used the native resolution of the projector (912 x1140 pixels), which illuminated the entire walking arena with pixels of size 292 µm (along wind axis) x 292 (perpendicular to wind axis) µm. The majority of our experiments used a 60 Hz stimulus update rate; the exception is the glider experiments (Supplementary Fig. 7d), for which we used a 180 Hz update rate to get faster updating stimuli. The average intensity of the red light within the walking arena was 4.25 µW/mm^2^. Though all data presented in this article used blind flies, initial exploratory experiments used flies that were not blind. To remove visual effects from the stimulating red light, we shone green light using an LED (Luxeon Rebel LED 530 nm) throughout the arena to flood the visual response. Though this was not necessary for blind flies, we retained the green light throughout the experiments presented here to compare to past data.

The projector and camera have distinct coordinate axes – camera and projector pixels are different sizes and their native coordinates systems are not even the same handedness. To infer the virtual perceived stimuli for navigating flies, the transformation between a 2D camera coordinate **x**_cam_ and a 2D stimulus coordinate **x**_stim_. We assume that the two are related by a combination of linear transformations and translations:

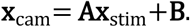

To estimate the matrix **A** and vector **B**, 3 mm diameter dots were projected at random locations 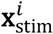 in the arena while recording with the camera; camera coordinates 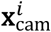 were determined in the imaged frame using the SimpleBlobDetector function in OpenCV. The 6 elements of **A** and **B** were then determined by minimized the least squares difference:

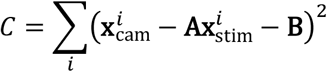

We verified manually that this procedure generated accurate transformations. We generated all stimuli using custom-written scripts in Python 3.7.4, and delivered these stimuli to the projector using the Python package PyschoPy, version 2020.2.4.post1.

### Electrophysiology

Single sensillum recordings from *Drosophila* antennae were performed as described previously (Gorur-Shandilya et al., 2017). The recording electrode was inserted into a sensillum on the antenna of an immobilized fly and a reference electrode was placed in the eye. Electrical signals were amplified using an Ext-02F extracellular amplifier (NPI electronic instruments). The ab2 sensillum was identified by i) its size and location on the antenna, and ii) test pulses of Ethyl 3-HyrdoxyButyrate, to which the B neuron is very sensitive. Spikes from the A and B neurons in this sensillum were identified and sorted as described previously (Gorur-Shandilya et al., 2017), using a spike-sorting software package written in MATLAB (Mathworks, Inc.) (https://github.com/emonetlab/spikesort).

### Experimental protocol

Experiments were carried out between 9 and 12 AM. All videos were 1 minute long, unless otherwise noted. Flies numbering between 10 and 30 were aspirated into the arena and let to acclimate for 2 minutes before experiments began. Before all experiments, optogenetic activation was verified by presenting static fictive odor ribbons (as in Supplementary Fig. 2c) with laminar wind for 120 seconds, and ensuring that flies followed the ribbons upwind as a positive control. Unless otherwise noted, each experiment ran for 60 seconds, with 60 seconds in between experiments. Throughout, experiments were interleaved such that the directions of the moving stimuli were randomized. No more than 30 videos were recorded on a single set of flies.

### Quantification of fly behavior and perceived fictive odor stimulus

#### Extraction of fly position, speed, and orientation from videos

All scripts were written in Python 3.7.4. Fly centroids were determined using SimpleBlobDetector in OpenCV, assuming a minimum area of 5 mm^2^. Given the centroids, fly identities were determined using custom tracking scripts. Briefly, centroids in subsequent frames were matched to the nearest centroid, and if the centroids could not be matched, they were marked as disappeared. Flies marked as disappeared for more than 30 frames (0.5 seconds) were then deregistered. Subsequent detected centroids were then marked as new fly tracks. Fly orientations *θ* were determined by first using the *canny* function in the Python module *scikit-image* to determine the points defining the fly edges around the centroid, then fitting these to an ellipse using custom-written Python scripts. Fly orientations are defined on the interval [0, 360°], but ellipse-fitting does not distinguish head (0°) from rear (180°). We properly resolved this using the fly velocity (below).

The above data defines the fly positions (*x, y*) and orientations *θ*. To remove measurement noise, we filtered each of these quantities with a Savitsky-Golay filter using a 4^th^-order polynomial and window size of 21 points (to avoid branch cuts in *θ*, it was first converted to an un-modded quantity). Velocities 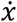 and 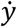 and angular velocity 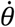 were defined by taking the analytical derivative of the fitted Savitsty-Golay polynomials for *x, y*, and *θ*. To resolve the two-fold symmetry in the fitted ellipses, and therefore distinguish the fly head from the rear, we used the fly velocity. For fly speeds greater than a given speed threshold, we matched the orientation to the fly velocity vector since flies walked forward. For other times, we matched the fly heading at the beginning and end of bouts when fly speed was below the speed threshold. The result was an estimate that may still have errors which occur as unnatural jumps in orientation. We repeated this process for various speed thresholds from 1 to 4 mm/s, and chose the orientation trace with the least number of jumps. We verified manually with several tracks that this procedure was highly reliable.

We noticed that during the experiments, particularly those with long fictive odor encounters such as the wide bars in Figs. 2 and 3, there was a slow, gradual bias toward one side of the arena (along the shorter axis of the arena). This only occurred for optogenetically active flies, and we reasoned it was due to a shadowing effect of the projector light from one antenna onto the other, since the projector lens is nearer to the bottom of its projected image. This shadowing effect essentially creates a static fictive odor gradient across the antenna. To account for this bias, we repeated all experiments that had an asymmetry in the perpendicular direction, such as bars perpendicular to the wind (Fig. 3d; 3^rd^ row), in both directions. We then averaged the turning biases from these two directions, after flipping the orientations appropriately. This would retain the effects due to direction sensing but remove the bias, under the assumption that this bias was an additive effect.

#### Estimation of perceived fictive odor stimulus in antennae

Given these smoothed and corrected *x, y, θ*, we then estimated the perceived fictive odor signal in the antenna region by defining a virtual antenna at a location 1.5 mm from its centroid along the ellipse major axis toward the fly head. To generate stable estimates – i.e. not relying on a single pixel value – we use the stimulus value averaged over a box of 0.25 mm^2^ around this location. Stimulus values in the antennal region are not measured by imaging, since the images are IR-pass filtered. Rather, they are obtained from knowledge of the stimulus pattern and the stimulus-to-camera coordinate transformation defined above. In PsychoPy, stimulus values are defined as 8-bit integers, from 0 to 255, but in practice we only deliver stimuli as max intensity (255) or 0. Accordingly, we treat the signal in the virtual antenna as binary, equal to 1 when the average stimulus value in the 0.25 mm^2^ region is above 200, and 0 otherwise.

#### Calculation of turning bias at bar edges

For the bar stimuli in Figs. 2-3, we identified ON and OFF edge hits as the times that the antennal signal switched from 0 to 1 or 1 to 0, respectively, where this binarization was calculated as described above. Correlated noise and glider stimuli (Fig. 4) were presented in blocks of 4s stimulus interleaved with 4s of no stimulus; thus the stimulus ON times were 0, 8, 16 seconds, etc. To calculated turning biases, we followed prior work and considered saccadic turning events, identified as points at which the absolute value of the angular velocity exceeded 100°/s, and ignored small jitters. Turn biases at a given time *t*_*i*_ (e.g. at an ON or OFF edge hit (Fig. 2-3)), were defined as the sign of the change in fly orientation from *t*_*i*_ + 150 ms to *t*_*i*_ + 300 ms, provided the absolute value of angular velocity in that window exceeded 100°/s at some point in that window. We used this 150 ms latency after *t*_*i*_ to account for uncertainties in *t*_*i*_ due to uncertainties in exact position of the antenna, which we estimated as being upper bounded by 2 mm. For correlated noise and glider stimuli, we considered orientation changes from *t*_*i*_ to *t*_*i*_ + 300 ms; the 150 ms latency was not needed in this case since the signal was independent of fly behavior, so the hit time was known to the precision of the inverse frame rate (16 ms). For all plots, to remove tracks in which flies may have been turning before the hit, we ignored points for which the absolute angular velocity exceeded 100°/s between 300 ms and 150 ms before the hit.

### Plume simulations

Direct numerical simulations were generated using the CFX® hydrodynamic simulation software package of ANSYS 2019. Parameters were chosen to emulate the flow and intermittent odor structure of the plume analyzed in Fig. 1 (Demir et al., 2020). An odorant with molecular diffusivity *D*_*m*_ = 7.3*e*-6 m^2^/s was injected mid-stream (vertically and horizontally). The odorant was modeled as a conservative, neutrally buoyant tracer. The dimensions of the computational model domain were 30×18×1 cm, approximately matching those of the walking arena (Demir et al., 2020). The computational air inlet boundary was modeled as a uniform velocity condition, representing an idealized collimated flow. The outlet boundary condition was modeled as a zero-pressure gradient opening allowing for bidirectional flow across the boundary. Walls were modeled using hydraulically smooth, no-slip boundary conditions. To reproduce the stochastic airjets creating the complex flow and plume, alternating jet pulses of air were applied from two orifices on opposite sides of the flume. The time series of pulses were identical to the experiments (Demir et al., 2020). The model domain was broken up into 4.7*e*6 tetrahedral elements where velocity and concentration were computed, with the largest element’s length at 5 mm with an inflation layer along the domain boundaries and a refined mesh around the inlet orifices.

The flow was simulated at a 2.5 ms time step using a *k*-*ϵ* eddy viscosity model (Pope, 2011), which solves the Reynold-averaged Navier Stokes equations, where the momentum equation is defined as:

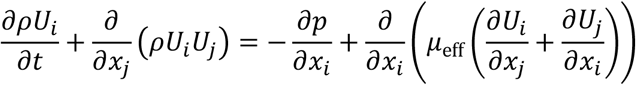

and the continuity equation as:

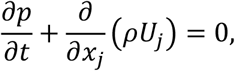

where *ρ* is the fluid density, *p* is pressure and *µ*_eff_ is the effective fluid viscosity. The turbulent eddy viscosity is treated analogously to viscosity in laminar flow such that *µ*_eff_ = *µ*_*t*_ + *µ* where *µ*_*t*_ is the turbulent viscosity and *µ* the fluid viscosity. The *k*-*ϵ* model assumes the local turbulent viscosity is related to the local turbulent kinetic energy (*k*) and the eddy dissipation rate (ε) as follows:

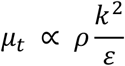

The advection-diffusion equation for conservative tracers was used to model the chemical transport of the odorant:

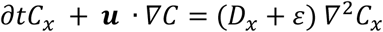

where *C*_*x*_ is the tracer concentration, ***u*** is the velocity field, *D*_*x*_ is the molecular diffusivity and *ε* is the local eddy diffusivity solved from the turbulence model. For all further analysis, we used the concentration and velocity in a plane 1 mm above the bottom of the domain, in the approximate z-plane of the fly antennae.

### Mathematical modeling and data analysis

#### Inter-antennal latency of edge hit Δ*T*

The inter-antennal latency Δ*T* as a function of fly walking speed |**v**_fly_| and bar speed |**v**_bar_| can be calculated with basic geometric considerations. Here, we assume that the fly speed along the bar direction is sufficiently slow such that the bar passes over the fly. Consider a coordinate system in the frame of the moving bar, where the bar direction is +*y* (i.e. the bar’s edge is in *x*). The fly velocity in this frame is

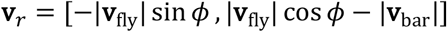

where *ϕ* is the angle of rotation from **v**_bar_ to **v**_fly_ in the experimenter frame. The inter-antennal latency Δ*T* is then the projection of the antennal spacing *L* along **v**_bar_ divided by the projection of **v**_*r*_ along **v**_bar_. The former is *L* sin *ϕ* and the latter is the *y*-component of **v**_*r*_; the sign of *L* sin *ϕ* is treated as meaningful, so that a positive/negative value means the left/right antenna is hit first. Thus:

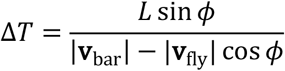

where the sign is given by the numerator since the denominator is always positive for bars passing over the fly.

This expression ignores the fly’s angular velocity while walking. Assuming that the fly is walking forward while also turning at a rate *ω*, then the total accumulation of orientation over the Δ*T* interval is *ω*Δ*T*, which for typical values of the maximum rotation rate during normal turns *ω* ∼ 300°/s and typical inter-antennal latencies without turning, Δ*T* < 15 ms, is less than 5 degrees. This would be if the fly were turning at a maximum angular velocity. For more typical jitters, rotation rates are approximately 20°/s (Demir et al., 2020), giving an accumulated angle during of less than 1 degree. If we incorporate this error as an uncertainty on *ϕ*, δ*ϕ*, then Δ*T* acquires an error of

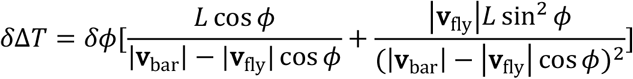

With the values assumed throughout, |*δ*Δ*T*| < 1 ms, so *ω* is safely ignored to the resolution of our experiments.

#### HRC output versus Δ*T* for traveling edges

Our prediction for the turning bias as a function of the latency Δ*T* at which an edge of odor hits the right antenna after hitting the left, is based on the output *r*(*t*) of the mirror-symmetrized Hassenstein-Reichardt correlator (Salazar-Gatzimas et al., 2016). To calculate *r*(*t*), we model the correlator architecture as depicted in Fig. 4a. Specifically, the time-varying signals from the 2 sensors are *s*_*L*_ (*t*) and *s*_*R*_ (*t*). In one arm of the computation, *s*_*L*_(*t*) is linearly filtered with an exponential 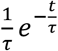, while *s*_*R*_(*t*) is transmitted unchanged; these are then multiplied. For a traveling ON edge moving left to right, we have *s*_*L*_(*t*) = *H*(*t*) and *s*_*R*_ (*t*) = *H*(*t* − Δ*T*), where *H*(·) is the Heaviside function. Then the product of the filtered values is:

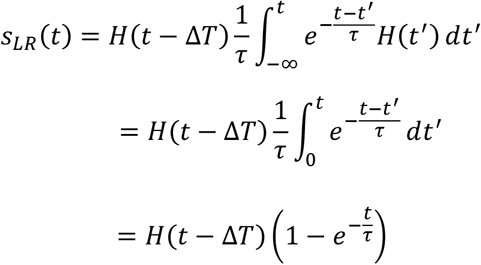

The other arm is similar, except that *s*_2_(*t*) is filtered and *s*_1_(*t*) is transmitted unchanged. Then the product of the filtered inputs is:

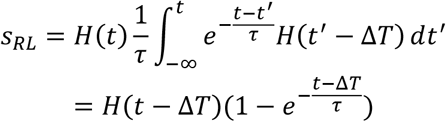

The correlator output is therefore:

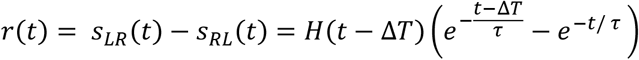

Assuming that flies sense odor direction using this computation, the output of the correlator, *r*(*t*), must be converted to a behavior; here, we model this behavior as the turning bias being proportional to ∫ *r*(*t*)*dt*:

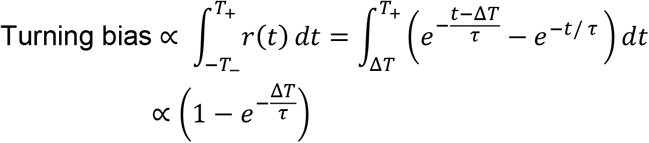

provided that behavioral timescales *T*_−_ and *T*_+_, over which the correlator response is integrated to produce the turning response, are large compared to *τ* and to Δ*T*. Long after the edge hit, *t* ≫ *T*_−_, the signals are both *s*_*L*_ = *s*_*R*_ = 1, giving an HRC output of 0, as expected for the anti-symmetric architecture.

To estimate the filtering constant *τ*, we minimize:

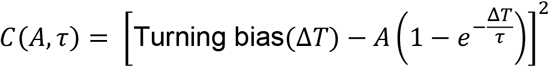

over *A, τ*. The turning bias is plotted in increments of Δ*T* = 4 ms, where the value at a given Δ*T* includes values from ± 4 ms. Neighboring points therefore contain overlapping data; this has the effect of smoothing – but not biasing – the turning bias vs. Δ*T* curve.

Responses to rightward moving OFF edges are analogous. The signal switches from 1 to 0 at the OFF edge (set it to *t* = 0), so the signal on the left sensor is *s*_*L*_(*t*) = 1 − *H*(*t*) and for the right sensor is *s*_*R*_ = 1 − *H*(*t* − Δ*T*). Then one arm of the HRC is:

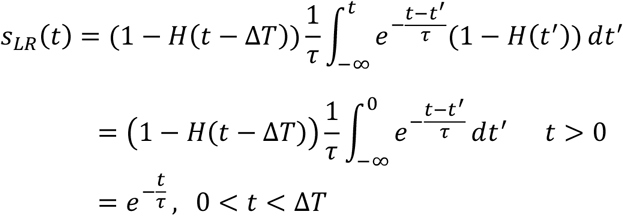

and *s*_*LR*_(*t*) = 0 for *t* > Δ*T* and *s*_*LR*_(*t*) = 1 for *t* < 0. The other arm output is simply *s*_*RL*_ = 1 for *t* < 0 and *s*_*RL*_ = 0 for *t* > 0, since the non-delayed arm drops to zero as soon as the edge passes it at *t* = 0. Thus the output is:

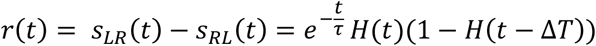

Integrating this quantity over time gives the same turning bias as the ON edge.

#### Generation of correlated noise stimuli and *C*(Δ*x*, Δ*t*)

Correlated noise stimuli were generated as previously described (Salazar-Gatzimas et al., 2016). We used optogenetic bars that were parallel to the short axis (*y*) of the arena (e.g. perpendicular to the wind direction, which runs along *x*). Each bar has a width of one *x*-pixel – thus, refer to an *x*-pixel as a “pixel,” since correlations are defined just in the *x*-direction. The stimulus value (where -1 and 1 are for dark and bright bars, respectively) of a bar at pixel location *x* and time *t* is given by *c*(*x, t*) = sgn(*η*(*x, t*) + *αη*(*x* + *β*Δ*x, t* + Δ*t*)), where each *η*(*x, t*) is independently chosen from a standard normal distribution. Δ*x* is the pixel spacing; Δ*t* is the inter-frame interval. The constant *β* governs the direction of the correlations: +1 for stimuli correlated in the +*x* direction (“with-wind” in the main text) and -1 for stimuli correlated in the −*x* direction (“against-wind”). The constant *α* governs the polarity of the correlations; +1 or −1 for positive or negative correlations, respectively.

The correlations can be computed straightforwardly (Salazar-Gatzimas et al., 2016). Assume that *α* = *β* = 1; the other cases are analogous. The correlations between two pixels separated by spacing *x*′ and timing *t*′ we denote *C*(*x*′, *t*′) = ⟨*c*(*x, t*)*c*(*x* + *x*′, *t* + *t*′)⟩. In general,

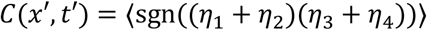

where *η*_*i*_ is one sample of *η*. For most choices of *t*′, *x*′, all *η*_*i*_ are distinct, so the correlation reduces to 0 since the sums are independent. For *x*′ = *t*′ = 0, the correlation reduces to the variance of *c*(*x, t*), which is 1. However, for *t*′ = Δ*t* and *x* ′ = Δ*x, η*_2_ = *η*_3_. Then,

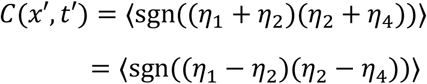

since the random variables are symmetric about 0. The sign depends only on the ordering of the *η*_*i*_, which are 3 independent samples from a standard normal distribution. There are 6 ways to uniquely order the *η*_*i*_, only two of which give a positive sign (*η*_1_ > *η*_2_ > *η*_4_ and *η*_1_ < *η*_2_ < *η*_4_); thus the expected value is 1/3 (Salazar-Gatzimas et al., 2016). An analogous property holds for *t*′ = −Δ*t,x*′ = −Δ*x*. Finally, the *α* and *β* factors are incorporated straightforwardly as scale factors, giving:

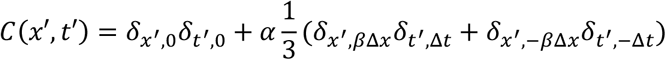

Note that the correlation can be calculated by averaging over all of spacetime, or just in space for a fixed set of times, or just in time for a fixed set of points. The latter is our interpretation for the HRC output from fixed antennae, assuming the correlation direction is perpendicular to the fly body.

#### Generation of glider stimuli

Here, the stimulus value of a bar at pixel location *x* and time *t* is given by *c*(*x, t*) = *B*(*x* − *βt*Δ*x*/ Δ*t*), where *B* = 2*X* − 1 with *X* ∼ Bernoulli(*p* = 0.5), Δ*x* is the pixel spacing, and Δ*t* is the inter-frame interval. The correlation between two pixels separated by spacing *x*′ and timing *t*′ is

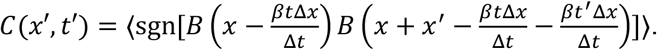

Then, *C*(*x*′, *t*′) = 1 when 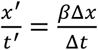– i.e., the correlation matrix has a diagonal or antidiagonal structure for *β* = 1 and *β* = −1, respectively. These stimuli are a class of *glider* stimuli with a two-point correlation structure. Visually, these gliders are a frozen pattern of random light dark bars moving statically at constant speed in the *βx* direction.

#### HRC output for correlated noise stimuli

Here we calculate the HRC output for correlated noise stimuli. Assume that the antennae are held at approximately the spacing of the correlation shift Δ*x* (see last section), and that the correlation direction is +*x* (rightward over the fly body), so *β* = 1 from the last section. Then one arm of the HRC gives:

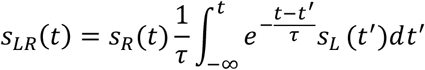

Averaging over time gives:

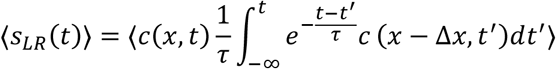

Since *β* = 1, then only the last term in the correlation equation applies:

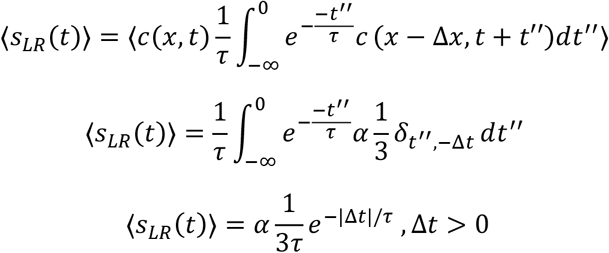

This equation holds for Δ*t* being positive. The other arm is analogous, for Δ*t* < 0.

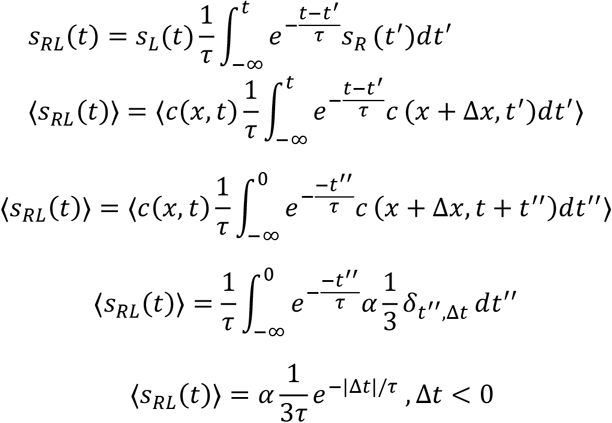

Thus, the full correlator output is

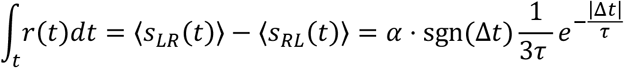

Note that the correlator output response switches sign if the correlation polarity *α* flips – this is the reverse phi response. There is a slight artificiality in this expression, in that the response is discontinuous at Δ*t* = 0. We have assumed an exponential filter, which technically has an immediate response time, violating causality. In addition, the optimal response occurs for an inter-frame interval Δ*t* that is arbitrarily small. As a more realistic filter, one can use 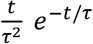, which has zero response at time zero and maximal response at *t* = *τ*. Then:

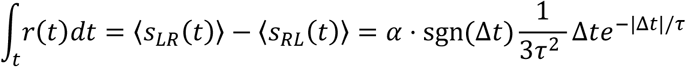

This filter is continuous at Δ*t* = 0, and the maximum correlator output occurs when the filter timescale *τ* matches the interframe interval Δ*t*. In either case, the salient point is that the response is antisymmetric in both the temporal shift Δ*t* and the correlation polarity *α*, as expected.

#### Analysis of imaged plume

We re-analyzed behavioral data previously extracted from *Drosophila* navigating an imaged complex plume of smoke (Demir et al., 2020) in the same walking assay used throughout this study. The signal in the virtual antenna was quantified as described previously; briefly, the virtual antenna is defined as an ellipse perpendicular to the body axis with the long axis given by the size of the fly (1.72 ± 0.24 mm) and the small axis equal to one-fifth the minor axis of the fly (0.46 ± 0.24 mm). We re-analyzed the imaged fly and signal data to resolve the virtual antenna signal into 14 pixels along its long axis (averaged along its short axis). Thus, the signal is a vector ***s***_ant_(*t*) = [*s*(*x*_1_, *t*), *s*(*x*_2_, *t*), …, *s*(*x*_14_, *t*)] defined at locations along the antenna’s long axis ***x***_ant_ = [*x*_1_, …, *x*_14_] for a given time *t*.

The overall concentration in the antenna was calculated as the average signal over the center of the virtual antenna – at the locations [*x*_5_, *x*_6_, *x*_7_, *x*_8_]. The gradient ∇*c*_ant_ in the virtual antenna at a given *t* was calculated by regressing ***s***_ant_ against ***x***_ant_ and extracting the slope. The odor velocity in the virtual antenna was estimated by calculating correlations of the virtual antenna signal over space and time. For a given *t*, we calculated 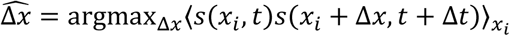, where Δ*x* spanned integers from -7 to 7, and Δ*t* is the interframe interval (11 ms), and *s*(·) were mean subtracted. This gives the signed number of pixels for which the correlation between two successive frames is maximized, up to the length of the antenna. The odor velocity was then defined as 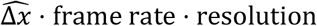, where the frame rate is 90 frames per second and the spatial resolution is 0.153 mm per pixel. We disregarded points for which 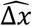 was ±7, since those may not represent local maxima but were instead limited by the size of the antenna. All three quantities – total concentration, gradient, and odor velocity – were smoothed in time using a Savitsky-Golay filter of order 2 and smoothing window of 25 timepoints ∼ 270 µs.

To remove boundary effects from the arena extent, we only used for Fig. 1c-e points for which the fly was in the central region of the arena, 100 < *x* < 250 mm, |*y* − *y*_0_| < 40 mm, where *y*_0_ is the plume’s central axis, and only points for which fly speed was greater than 0.1 mm/s. Angular velocity was calculated as the average orientation change over 200 ms.

#### Analysis of simulated plume

The simulation generated concentration fields *c*(*x*_*i*_, *y*_*i*_, *t*) and flow velocity fields **v**_wind_(*x*_*i*_, *y*_*i*_, *t*) defined on grid points (*x*_*i*_, *y*_*i*_) of a non-uniform mesh. We first generated values on a 0.5 mm square lattice, by triangulating the data and performing barycentric linear interpolation over each triangle (*scipy*.*interpolate*.*griddata* in Python, with method ‘*linear*’). Fields in Fig. 5 and Supplementary Fig. 8 were plotted every 1 cm, (i.e. every 20 pixels on the original 0.5 mm lattice). Wind speed vectors at each point on this 1 cm lattice were generated by averaging **v**_wind_ over the 20 × 20 values in a 1 cm^2^ box. The plotted **v**_wind|odor_ field was generated by only considering wind vectors for which the odor concentration was above 1e-3. Odor gradients were generated by calculating local differences ∇*c*_*x*_ and ∇*c*_*y*_ in the *x*-and *y*-directions, respectively. Specifically, for ∇*c*_*x*_, we calculated (*x*_+_ − *x*_−_)/ (*x*_+_ + *x*_−_), where *x*_+_ and *x*_−_ were the averages in the right and left half of a 1 cm^2^ box centered at each lattice point, respectively. ∇*c*_*y*_ was calculated analogously, using the top and bottom half of the same box. Odor velocities were calculated similarly to those in the imaged plume used in Fig. 1, by correlating the values in a given spatial region between two frames. Specifically, to get **v**_*x*, odor_ at a given time *t*, we calculated 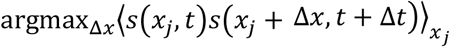, where *s*(*x*_*j*_, *t*) was the odor concentration in a 1 cm^2^ box averaged over the *y*-direction for each *x*_*j*_ pixel spaced by 0.5 mm. The shifts Δ*x* ran from -20 to 20 pixels (±1 cm). This quantity was multiplied by the frame rate 100 frames per second and by the spatial resolution 0.5 mm per pixel to get **v**_*x*, odor_ in mm/s. An analogous operation was done for **v**_*y*, odor_ using the same 1 cm^2^ box. All odor gradient and odor velocity values for very low odor concentrations were set to Nan, as were any odor velocity values that produced a maximum shift |Δ*x*| = 20. The resulting wind speed, gradient, and odor velocity were all smoothed in time using a Savitsky-Golay filter of order 1 and window length 11 (110 ms).

#### *In silico* virtual agent model and simulation

Virtual agents with 2 spatially separated sensors navigated the simulated plume described above using a simple algorithm. All agents were initialized at the back of the arena, facing upwind. At each frame (10 ms), agents turned either left or right 90° (except in one case where they maintained their heading; see below), depending on the navigation strategy as described in the main text, and stepped forward 0.75 mm. The sensors were placed 0.5 mm to the left or right of the agent centroid. The measured odor signal concentration was defined as 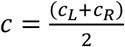, where the concentration in each sensor was *c*_*L*_ and *c*_*R*_, respectively. We set the detection threshold at *c*_0_ =1e-3. The odor correlation between the two sensors was defined as *c*_odor_(*t*) = *c*_*L*_ (*t*)*c*_*R*_ (*t* + Δ*T*) − *c*_*L*_ (*t* + Δ*T*)*c*_*R*_ (*t*), where the delay timescale Δ*T* was chosen as 1 frame. From *c*_odor_, the odor direction *v*_odor_ was defined +1 if abs(*c*_odor_(*t*)) > 1e-8 and sgn(*c*_odor_(*t*)) > 0, as -1 if abs(*c*_odor_(*t*)) > 1e-8 and sgn(*c*_odor_(*t*)) < 0, and as 0 otherwise. In general, odor signals with a leftward component over the virtual agent in its body frame had *v*_odor_ = 1 and, while those with a rightward component had *v*_odor_ = −1. Simulations were carried out separately for agents that could sense (DS+) and could not sense (DS-) odor direction. Agents followed the strategy as described in the main text. For DS+ flies, whenever *c*_odor_ was below threshold (abs(*c*_odor_(*t*)) >1e-8), but the odor was still detectable (*c* > *c*_0_), the decisions obeyed the DS-strategy.

#### Theoretical analysis of odor motion in turbulent odor plumes

Here we investigate the motion of odor signals perpendicular to the mean flow using a toy model of turbulent plume similar in spirit to those used in (Balkovsky and Shraiman, 2002; Goldstein, 1951; Taylor, 1922). Odor packets are released from a point source at a given rate. The concentration around the center of each packet is given by a local diffusive process that spreads the concentration via molecular diffusion of the odor. Meanwhile, the packets themselves are advected downwind by the mean flow, while being dispersed by the fluctuating velocity *u* (Taylor, 1922). We consider the simple case of an isolated packet and calculate its expected velocity crosswind to the flow, at different locations throughout the plume. For analytical simplicity, we model the turbulent velocity *u* as a telegraph process that switches between left motion and right motion at speed *v*, where the switching rates from left to right and vice versa are both *λ* = 1/ *T*. Thus, 2*T* is equivalent to the Lagrangian integral time scale and the packet speed *v* to the r.m.s. of the turbulent velocity field. While the velocity *u* switches discontinuously between +*v* and −*v*, its time correlation function is the same as that of the Ornstein-Uhlenbeck (O-U) process often used to model homogeneous isotropic turbulence (Pope, 2011; Taylor, 1922):

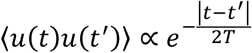

Our goal is an estimate of the average odor motion velocity at a given lateral distance from the plume, at a given time *t*, ⟨*v*⟩_*y,t*_. Since packets are advected downwind at some speed *U* ≫ *v*, we have *t* ≈ *x*/ *U*, so that this is equivalent to finding the average lateral velocity at some *x, y* position in the plume (Pope, 2011). Run times are distributed as 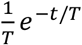, so packets reaching a given *y* will have been traveling for some distance 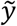, where 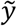 is distributed as 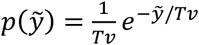. If the packets were originally uniformly distributed, then the average velocity at *y* would be 0. However, an asymmetry arises due to the non-uniform packet distribution, which is dispersing laterally from a delta function at *y* = 0. For times *t* ≫ *T*, the distribution of packets is approximately the diffusion kernel with effective turbulent diffusivity *D*_*T*_ = *Tv*^2^/ 2:

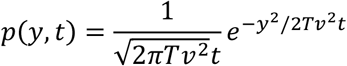

Under these assumptions, the average velocity at the fixed point ⟨*v*⟩_*y,t*_ is:

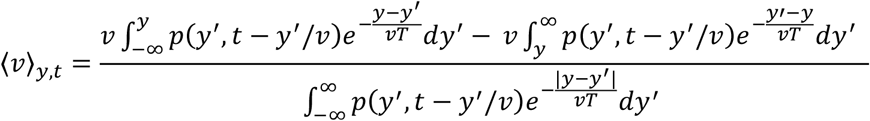

The first term in the numerator is for packets reaching *y* that have come from its left (these are traveling in the +*y* direction), while the second is for those reaching *y* that have come from the right, which are traveling in the −*y* direction. The denominator is a normalization factor given by the total number of packets reaching *y* at time *t*. This equation can be integrated numerically. To obtain an analytical approximation, we neglect the change in the packet distribution over the time of traveling one correlation time, approximating *p*(*y*′, *t* − *y*′/ *v*) by *p*(*y*′, *t*), since the packet distribution does not change appreciably over that time (the validity of this assumption was verified by simulations). Integrating:

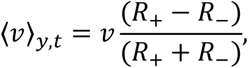

where

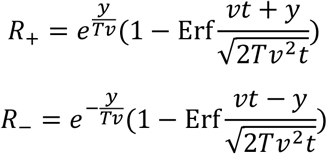

for |*y*| < *vt*, and 0 otherwise. We are interested in i) whether the average lateral velocity of the packets is directed outward from the plume, which would be indicated by an asymmetrical dependence in *y*, and ii) how this asymmetry depends on the correlation time *T*. The profile of ⟨*v*⟩_*y,t*_ is odd for all *T* (Supplementary Fig. 9a), indicating that for any *T*, the velocity of odor packets in the crosswind direction points away from the plume’s central axis. Moreover, for higher *T*, the velocity component points more strongly outward through a larger portion of the plume, indicating that correlations in the packet motion underlie this directional cue (Supplementary Fig. 9a).

We next investigate how the combination of packet diffusion and packet centroid motion together can influence a spacetime correlation of the odor concentration, as would be computed by time-resolved bilateral measurements. We define a lateral correlator ⟨Δ*y*Δ*t*|*y*_*i*_ ⟩ at a position *y* and time *t*, assuming a packet is traveling nearby with trajectory *y*_*i*_ (*t*). The correlator has the following form:

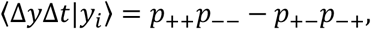

where

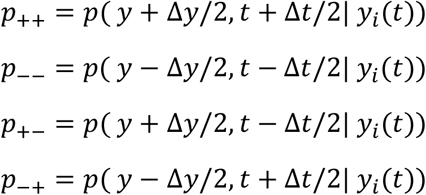

and where *y*_*i*_ (*t*) is the centroid of a nearby packet and *p*(·) is the local concentration at a given location and time around the packet. Thus, the correlator ⟨Δ*y*Δ*t*⟩ is a time-antisymmetrized quantity that compares the correlation of the odor concentration between two points in the direction perpendicular to the mean wind, separated by Δ*y* at times separated by Δ*t*, given a packet whose center is at (*x*_*i*_, *y*_*i*_) and which is released at *t* = 0. We stress that we do not imply that this correlator is being enacted by any circuitry, nor is it a unique definition. However, it has key features – namely comparisons across space and time, and time antisymmetry – which we will show to be sufficient to detect the lateral odor velocity. Expanding this correlator gives

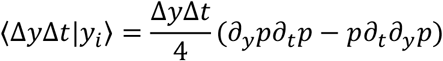

to lowest order. For the packet model, at appreciable times *t* ≫ *T*, this gives:

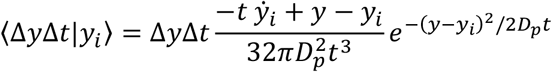

Note that this is for a single packet, and must be averaged over the packet distribution *p*(*y*_*i*_, *t*) to get the correlator at a fixed *y, t*:

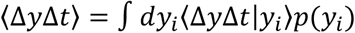

where 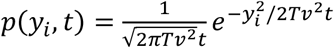 for *t* ≫ *T*, as above. We can approximate 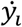 by 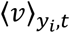 – the average velocity for a packet at position *y*_*i*_ as derived above. The expression for ⟨Δ*y*Δ*t*⟩ does not lend itself to a closed-form expression due to the complexity of 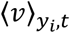; we integrate it numerically. We find that for *D*_*p*_ ≪ *D*_*T*_ = *v*^2^*T*/ 2, ⟨Δ*y* Δ*t*⟩ has a clear asymmetry about *y* = 0 as expected, and that the peaks are stronger with increasing correlation time *T* (Supplementary Fig. 9b). Moreover, ⟨Δ*y*Δ*t*⟩ increases on average with *v*, while decreasing with *D*_*p*_ (Supplementary Fig. 9c), indicating that the response essentially derives from correlated motion over the detector rather than molecular diffusion alone.

#### Statistical quantification

All error bars, when shown, represent standard error of the mean. Statistical tests used and significance levels (*p* value) for given comparisons are indicated in the main text. Throughout, *, **, ***, and **** refer to *p*-values of < 5e-2, <1e-2, <1e-3, and <1e-4. In some instances, **** may refer to *p* < 1e-6, if indicated in the text.

## SUPPLEMENTARY FIGURES

**Supplementary Figure 1.**
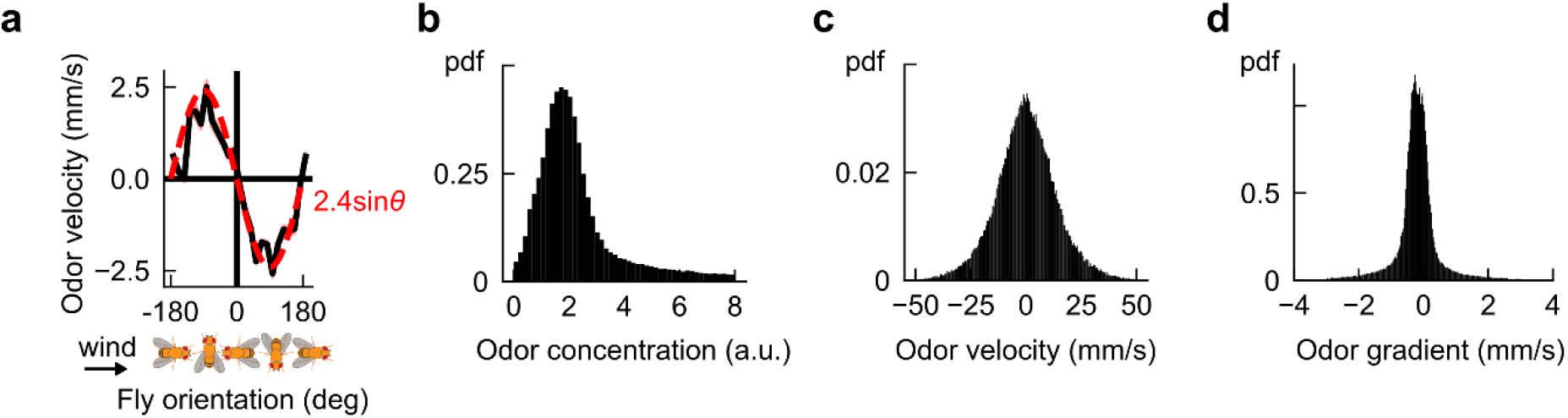
Verification of odor velocity calculation and distributions of signal-derived quantities in measured plume. **a**, Odor velocity measured in the virtual antenna at all times for navigating flies in measured smoke plume, plotted as a function of fly orientation. The sin(*θ*) trend reflects the fact that the main component of odor velocity is parallel to the mean wind direction 0°, as expected – a consistency check on the odor velocity calculation. **b**, Histograms of signal-derived quantities measured in the fly virtual antenna; the *x*-axis limits in Fig. 1c-e are determined by the extent of these histograms.

**Supplementary Figure 2.**
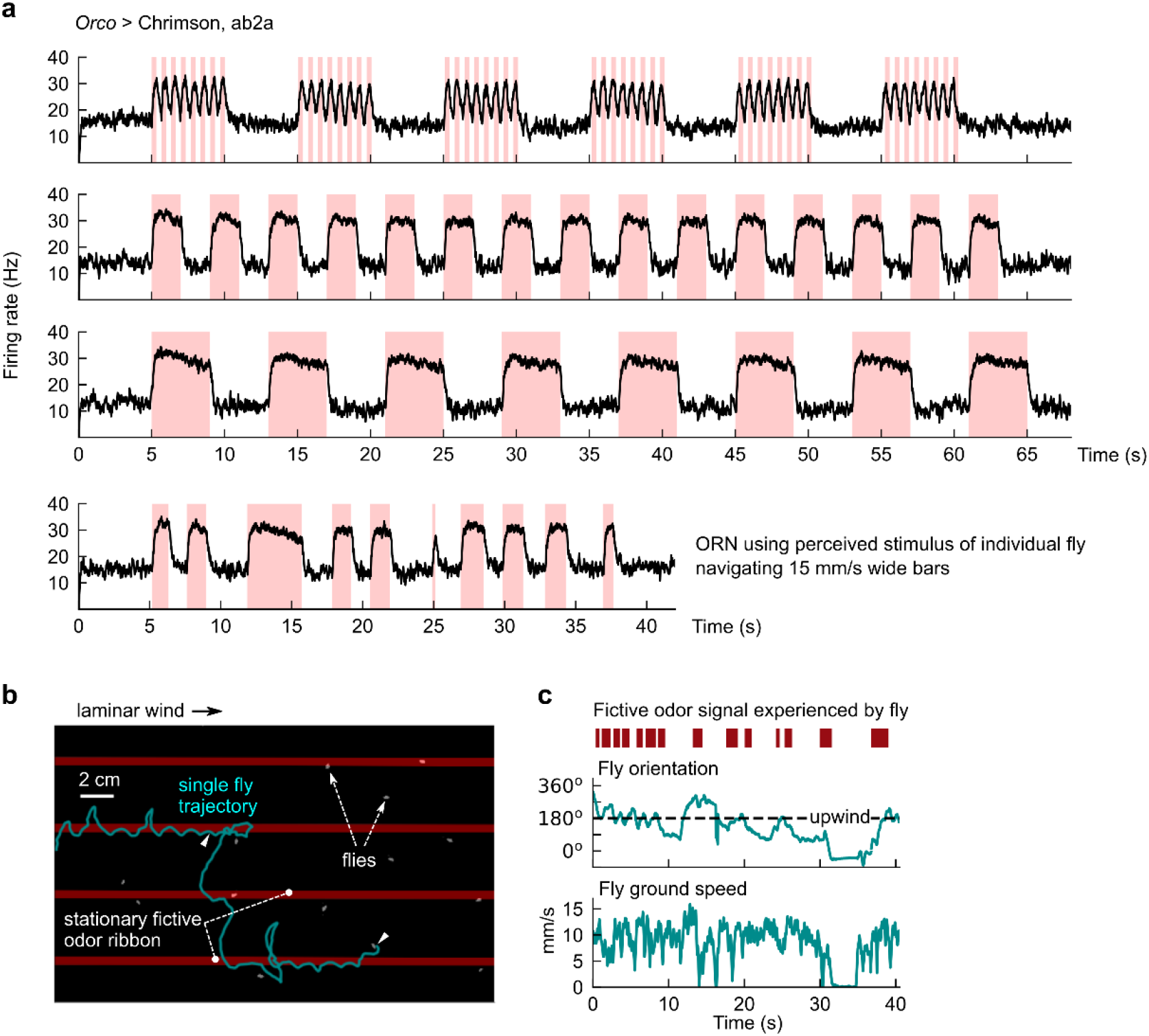
Electrophysiological and behavioral verification of optogenetic activation of *Drosophila* ORNs. **a**, Extracellular measurements of ab2A firing rates for various odor signals mimicking those we use throughout our study. Stimuli (red shades) are delivered using a Luxeon Rebel 627 nm red LED (Lumileds Holding B.V., Amsterdam, Netherlands) at 10 uW/mm^2^. The frequency and duty cycle for the stimuli in the first plot are 1.5 Hz and 50% respectively, which mimics what a stationary fly in the 5 cm wide, 15 mm/s fast moving bars (Fig. 2b) would perceive. Longer stimuli approximate the experienced stimuli in the wide moving bars (Fig. 2e-f). Last plot shows the perceived stimulus and corresponding firing rate for one representative measured fly navigating 15 mm/s moving wide bars. **b**, Illustrative track of fly following stationary fictive odor ribbons upwind. Red bars: optogenetic stimulus location – bars are overlaid on the figure, but not actually imaged since the image is IR-pass filtered. **c**, Perceived fictive odor signal for fly (red bars) can be simultaneously quantified with fly behavior (teal) by aligning camera and projector coordinate systems (Methods). Plotted are the perceived fictive odor signal and behaviors for the track shown in **b**.

**Supplementary Figure 3.**
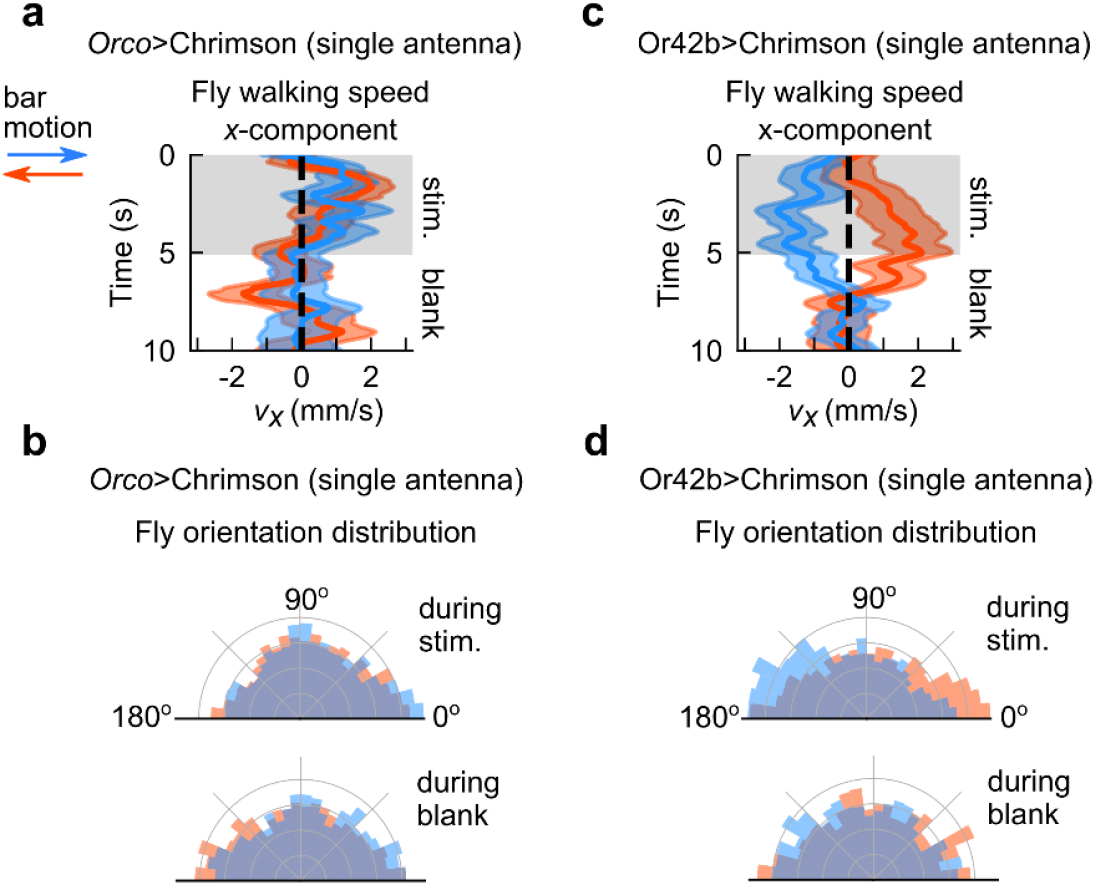
Olfactory direction selectivity is abolished in single antenna flies and preserved in flies expressing Chrimson in a single Or. **a**, Component of fly walking velocity along +*x* direction during the 5s stimulus (shaded grey) and blank periods (illustrated in Fig. 2b), in *Orco*>Chrimson flies who have one antenna ablated (compare to Fig. 2d). Blue and orange denote rightward and leftward moving bars, respectively. Since it is difficult to distinguish flies walking on the top and bottom surface of the assay, right- and left-antenna ablated flies are pooled. *n =* 307, 304 tracks for rightward and leftward bar motion, respectively. **b**, Distribution of fly orientations during the 5s stimulus (top) and 5s blank periods (bottom), for rightward (blue) and leftward (orange) bar motion, *Orco*>Chrimson flies with one antenna ablated (compare Fig. 2d). Orientations are symmetrized over the *x*-axis. **c-d**, Same as **a**-**b**, for *Or42b*>Chrimson flies (not antenna ablated). *n* = 80, 96 tracks for rightward and leftward bar motion, respectively.

**Supplementary Figure 4.**
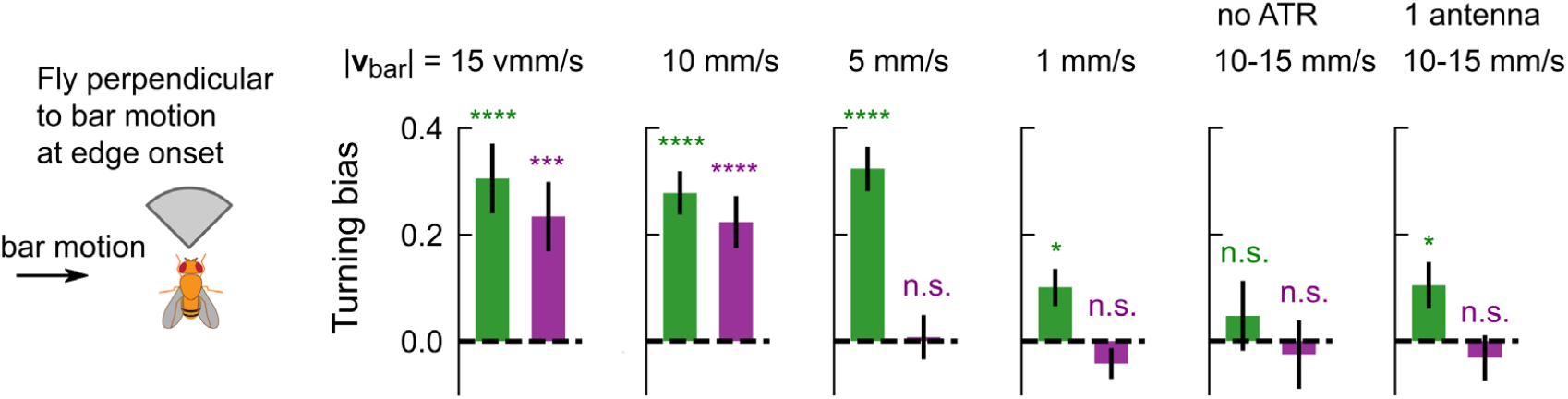
Turning responses at ON and OFF edges for moving bars at various speeds and negative controls are consistent with direction selectivity. Turning bias for all times that flies cross the fictive odor ON (green) or OFF (purple) edge, for flies oriented within a 90° sector of the direction perpendicular to bar motion. Turning bias calculated as sign of fly orientation change from 150 ms to 300 ms after the edge hit. All flies are *Orco*>Chrimson and fed ATR (i.e. optogenetically active) except in the 5^th^ plot, which are not fed ATR. Data are shown for bars that move at various speeds (left 4 plots), as well as for negative controls (5^th^ and 6^th^ plot). *P* values calculated using the chi-squared test (\*\*\*\**p* < 1e-4, \*\*\**p* < 1e-3, \*\**p* < 1e-2, \**p* < 0.05). *n* = 773, 1625, 1877, 1175, 3622, and 1487 tracks for the 6 plots, respectively). Direction selectivity is satisfied if both ON and OFF edge responses have the same sign; gradient sensing would require opposite signs for the two edges. Data indicate that flies counterturn against the direction of fictive odor bars at both edges, provided the bar speed is fast enough. Large ON responses for slow bar speeds are likely attributed to gradient sensing.

**Supplementary Figure 5.**
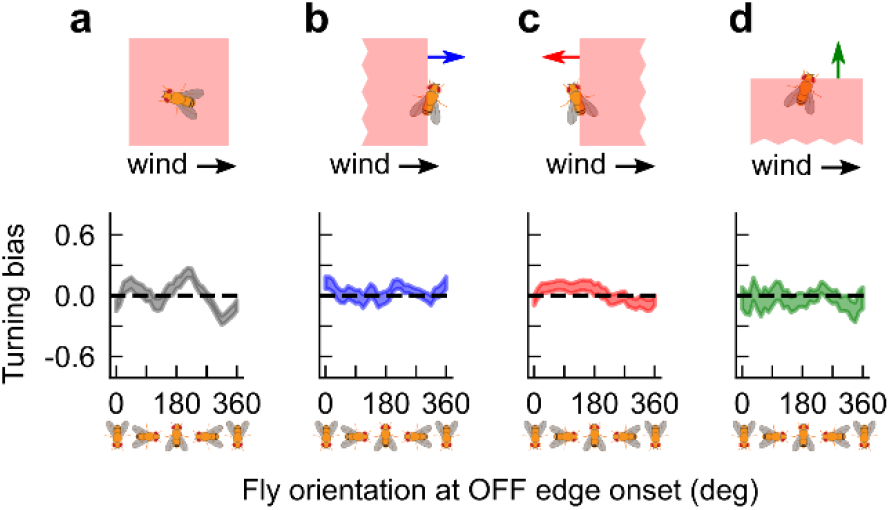
Fly turning to OFF edges in the presence of laminar wind exhibits no directional bias. **a**, Turning bias versus fly orientation when bilateral optogenetic stimulus is turned off (compare first plot in Fig. 3B for flash onset). **b**-**d**, Fly turning bias for 15 mm/s bars moving parallel, antiparallel, and perpendicular to 150 mm/s laminar wind (compare Fig. 3d).

**Supplementary Figure 6.**
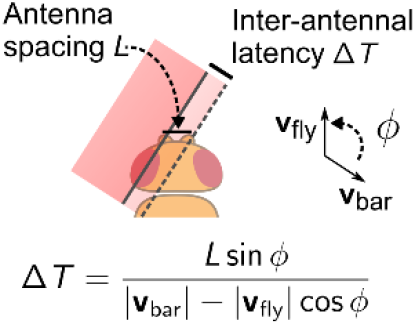
Schematic illustrating calculation of latency Δ*T* between antennae hits for moving edges. Correlation-based models for direction selectivity depend on the latency Δ*T* of the time the edge hits the two sensors – in this case, the fly’s two antennae. Measuring Δ*T* does not require resolving the image or stimulus at antennal resolution (∼300 *µ*m), rather Δ*T* can be inferred with knowledge of the fly’s orientation relative to the bar direction *ϕ*, as well as the speeds of the fly and bar – all of which are known. See Methods for details of the calculation and an estimate of the uncertainty.

**Supplementary Figure 7.**
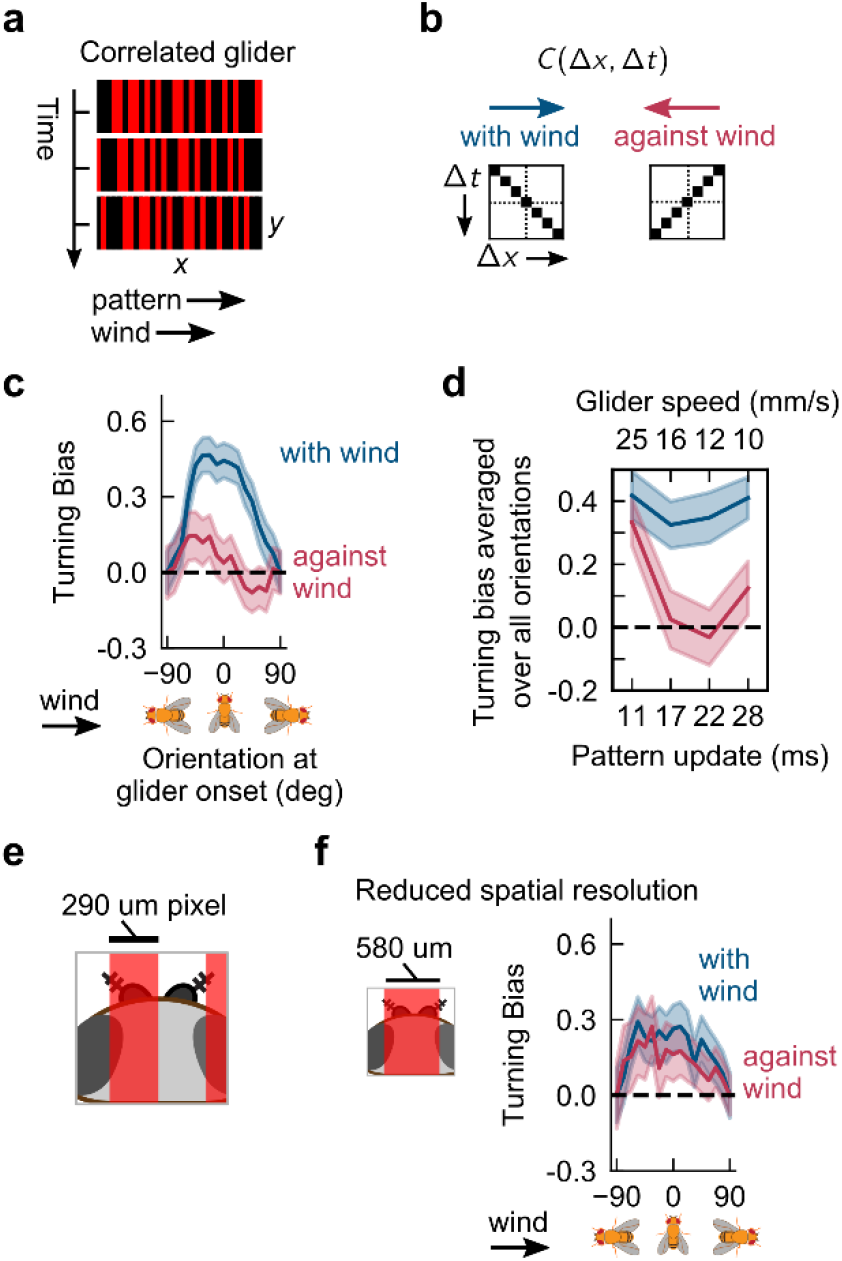
Gliders provide further evidence that direction sensing is enacted using a correlation-based algorithm. **a**, Snapshots of glider stimulus with correlations along +*x* axis, for 3 consecutive frames. In one instance of time, stimulus is a random pattern of light and dark 1-pixel-wide bars perpendicular to 150 mm/s laminar wind. Each *x*-pixel is perfectly correlated with the pixel to its right in the next frame; thus the pattern in the next frame is the same as the pattern in the current frame, but shifted by one pixel. Visually, this would be perceived as a fixed pattern moving coherently (“gliding”) to the right. **b**, Like correlated noise (Fig. 4 in main text), gliders are defined by their correlation matrix *C*(Δ*x*, Δ*t*). Unlike correlated noise, the correlations i) are exact – i.e. magnitude 1, and ii) exist for many spacetime points. That is, for rightward correlated gliders, a given pixel in a given frame is perfectly correlated with the pixel to its right one frame later, but also with the second pixel to its right 2 frames later, etc. Thus *C*(Δ*x*, Δ*t*) has values +1 along the diagonal. Similarly, *C*(Δ*x*, Δ*t*) has values 1 along the anti-diagonal. Since +*x* points downwind, we call gliders with correlations to the right “with-wind”, and gliders with correlations to the left “against-wind.” **c**, Turning bias versus fly orientation for with-wind (blue) and against-wind (red) gliders. Data using frame rates of 45 or 60 Hz are pooled. Gliders are presented in 4s blocks, interleaved with 4s of no stimulus. Turning bias is defined as the sign of the change in orientation from 200 to 500 ms after the block onset. We only used flies with speeds < 12 mm/s for gliders, since long-range correlations can interfere with the intended correlation if fly walking speed is near the glider speed. *n* = 597, 661 for with-wind and against-wind, respectively. **d**, Turning bias averaged over all orientations for different glider speeds. Glider speed is calculated as (pixel width)·(pattern update) where the pixel width is 290 µm and the pattern rate is some multiple of the inverse frame rate, 1/(180 Hz). *n =* 537, 289, 275, 440 tracks for with-wind stimuli at glider speeds 25, 16, 12, and 10 mm/s, respectively; *n* = 495, 308, 386, 383 tracks for against-wind stimuli at same glider speeds, respectively. **e**, For correlated stimuli to be perceived in our assay, the bar width (size of *x*-pixel, 290 µm), must be on the order of the fly antennal separation (∼300 µm). **f**, Glider stimuli experiments repeated for bars that were double the width, 580 µm. Differences now disappear for with and against-wind correlations, consistent with bilaterally-enabled direction sensing, since these bars are too wide to stimulate antennae differentially. *n* = 741, 677 for with-wind and against-wind, respectively.

**Supplementary Figure 8.**
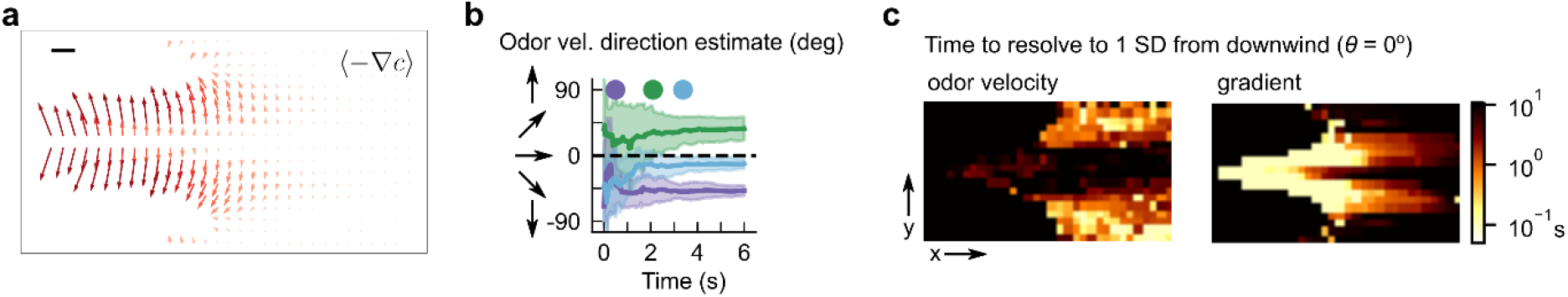
Odor velocity and concentration gradients provide complementary directional information in complex plumes. **a**, Vector field of the negative gradient of odor concentration −∇*c*, averaged over the full simulation (compare to Fig. 5c in the main text). Gradients contain strong lateral components near the odor source. **b**, Time course of an estimate of the direction of odor motion *θ*_odor_ = tan^−1^ (**v**_*y*, odor_, **v**_*x*, odor_) at the center of the boxed regions in Fig. 5a, determined by averaging all detectable *θ* in the past *t* seconds. Error bars are found by repeating this for 16 different 10 s time windows throughout the simulation, and taking the average and standard deviation over these 16 samples – these correspond to the mean and standard error of the mean. Dots indicate the time needed to distinguish the direction of odor motion from 0° (downwind) with a 68% confidence level for the 3 regions. **c**, Heatmap of time taken to distinguish the direction of odor motion from 0° to within 68% confidence for fixed locations throughout plume. Black values include the possibility that the odor motion direction is not distinguishable from downwind no matter how long one samples.

**Supplementary Figure 9.**
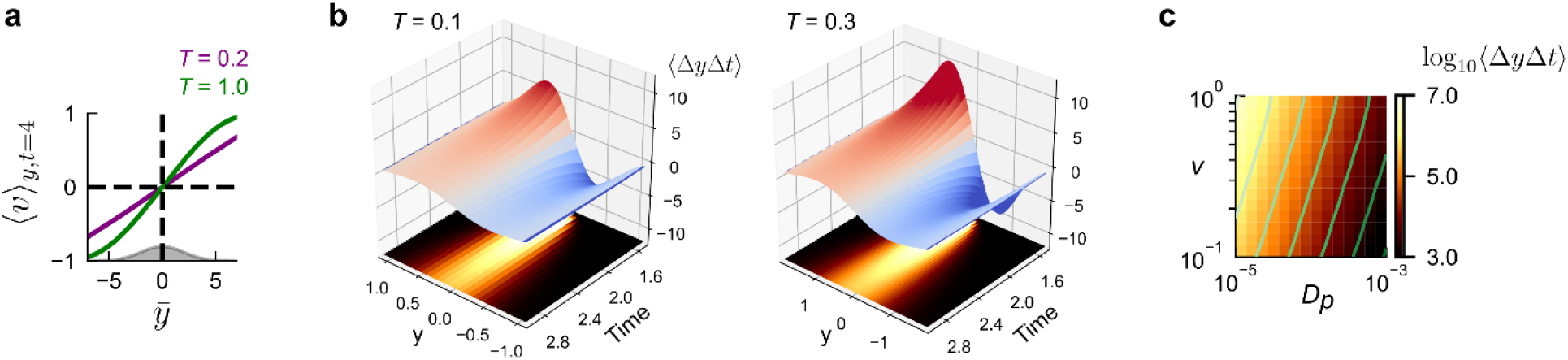
Odor velocity in model of turbulent plumes points outward from plume centerline and is computed by local space-time correlators. A packet model of turbulent plumes. Packets are released from a source and disperse in the lateral direction while being advected downwind (see Methods for model and calculation details). **a**, Packet velocity ⟨*v*⟩_*y,t*_ in the plume model, as a function of 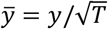, for two correlation times, *T* = 0.2 (purple) and *T* = 1 (green), at a fixed time *t* = 4. Here, *v* is set to 1. To directly compare velocity for plumes with different *T*, (and therefore different diffusivities) we plot the velocity versus the normalized length 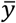. Specifically, since ⟨*y*^2^⟩ = 2*Tv*^2^*t* for *t* ≫ *T* then at a given *t*, the packet distribution in terms of 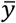 is the same for plumes with distinct *T*. The distribution of packets for either *T* is a function of 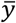 is shown in grey. The velocity is an odd function of *y*, i.e. it points outward from the plume axis. In addition, the asymmetry is steeper for higher correlation times. **b**, The value of the correlator ⟨Δ*y*Δ*t*⟩ as a function of lateral distance *y*, for various times *t* for *T* = 0.1 (left) and *T* = 0.3 (right). Here, *D*_*p*_ = 0.005. Since the packets are advected downwind with a velocity *U* ≫ *v*, then the time axis proportional to the downwind distance. The packet distribution is shown on the bottom; the limits of the *y*-axis are chosen such that the plume extents are the same in both plots. **c**, The total integral of the absolute value of ⟨Δ*y*Δ*t*⟩ at a fixed *t* = 4, as a function of odor packet speed (*y*-axis) and molecular diffusivity (*D*_*p*_), with *T* = 1, *v* = 1. The correlator is higher for greater packet speeds and lower molecular diffusivities (top left corner).

